# Perception of butenolides by *Bacillus subtilis* via the α/β-hydrolase RsbQ

**DOI:** 10.1101/2023.12.05.570042

**Authors:** Kim T Melville, Muhammad Kamran, Jiaren Yao, Marianne Costa, Madeleine Holland, Nicolas L Taylor, Georg Fritz, Gavin R Flematti, Mark T Waters

## Abstract

The regulation of behavioural and developmental decisions by small molecules is common to all domains of life. In plants, strigolactones and karrikins are butenolide growth regulators that influence several aspects of plant growth and development, as well as interactions with symbiotic fungi^1–3^. DWARF14 (D14) and KARRIKIN INSENSITIVE2 (KAI2) are homologous receptors that perceive strigolactones and karrikins, respectively, and that hydrolyse their ligands to effect signal transduction^4–7^. RsbQ, a homologue of D14 and KAI2 from the Gram-positive bacterium *Bacillus subtilis*, regulates growth responses to nutritional stress via the alternative transcription factor SigmaB (σ^B^)^8,9^. However, the molecular function of RsbQ is unknown. Here we show that RsbQ perceives butenolide compounds that are bioactive in plants. RsbQ is thermally destabilised by the synthetic strigolactone GR24 and its desmethyl butenolide equivalent dGR24. We show that, like D14 and KAI2, RsbQ is a functional butenolide hydrolase that undergoes covalent modification of the catalytic histidine residue. Exogenous application of both GR24 and dGR24 inhibited the endogenous signalling function of RsbQ *in vivo*, with dGR24 being 10-fold more potent. Application of dGR24 to *B. subtilis* phenocopied loss-of-function *rsbQ* mutations and led to a significant down-regulation of σ^B^-regulated transcripts. We also discovered that exogenous butenolides promoted the transition from planktonic to biofilm growth. Our results suggest that butenolides may serve as inter-kingdom signalling compounds between plants and bacteria to help shape rhizosphere communities.

## Results and Discussion

DWARF14 (D14) and KARRIKIN INSENSITIVE2 (KAI2) are paralogous α/β-hydrolases that function both as butenolide receptors and as enzymes that hydrolyse their respective ligands through a canonical catalytic triad of Ser, His and Asp residues^4,5,10,11^. Besides karrikins or their metabolites, KAI2 is likely a receptor for an unknown hormone provisionally known as KAI2-ligand (KL) that regulates seed germination, root colonisation by arbuscular mycorrhizal fungi, and tolerance to abiotic stress such as drought^12–16^. Close sequence homology of D14 and KAI2 to bacterial proteins, exemplified by RsbQ from *Bacillus subtilis*, suggests an ancient origin for this family of α/β-hydrolases by horizontal gene transfer to an algal ancestor of land plants^17–19^.

In *B. subtilis*, RsbQ is a positive regulator of the pathway that activates σ^B^, an alternative sigma factor that directs bacterial RNA polymerase to transcribe a suite of over 250 genes involved in stress response, environmental adaptation and biofilm formation^8,20–22^. RsbQ is encoded in a bicistronic operon with its signalling partner RsbP, a PP2C phosphatase. Loss-of-function *rsbQ* and *rsbP* mutants fail to induce expression of σ^B^-dependent transcripts in response to carbon limitation but respond normally to environmental stressors such as ethanol and salt^8,23,24^. Thus, RsbQ and RsbP define the “energy stress” branch of σ^B^ regulation associated with carbon limitation and the shift from logarithmic to stationary growth. However, the chemical signal that RsbQ and RsbP perceive to initiate the activation of σ^B^ is unknown.

RsbQ from *B. subtilis*, AtKAI2 from *Arabidopsis thaliana* and OsD14 from *Oryza sativa* are topologically highly similar^25–27^ (Figure 1A). All three have a characteristic cap domain that forms a hydrophobic pocket leading to a canonical catalytic triad. At least three of the pocket residues that are highly conserved among RsbQ homologues from Firmicutes (F27, Y125 and L144) are also common to the highly conserved clade of KAI2 proteins (eu-KAI2) in plants, but none are specific to RsbQ and D14 only (Figure 1B). The positions of these residues are highly correspondent in RsbQ and AtKAI2, and help to define a ligand binding pocket of RsbQ that is smaller than that of AtKAI2 and OsD14: the phenyl side chain of F197^RsbQ^ points towards the catalytic triad and partially occludes the pocket, whereas F194^AtKAI2^ and F245^OsD^14 are deflected away (Figure 1C–E). The similarities between RsbQ and KAI2 pockets may be functionally significant because Y124 has been implicated in ligand selectivity in KAI2, as have positions 157 and 161 ^28,29^, which are comparatively more variable among RsbQ proteins (Figure 1B). Such variable positions may reflect promiscuity in ligand selectivity between different RsbQ homologues, and/or the inherent evolutionary diversity within a large taxon such as Firmicutes. Regardless, given these close structural relationships, we hypothesised that RsbQ would mediate butenolide perception and response in *B. subtilis*, and with a ligand preference similar to KAI2 from plants.

**Figure 1.**
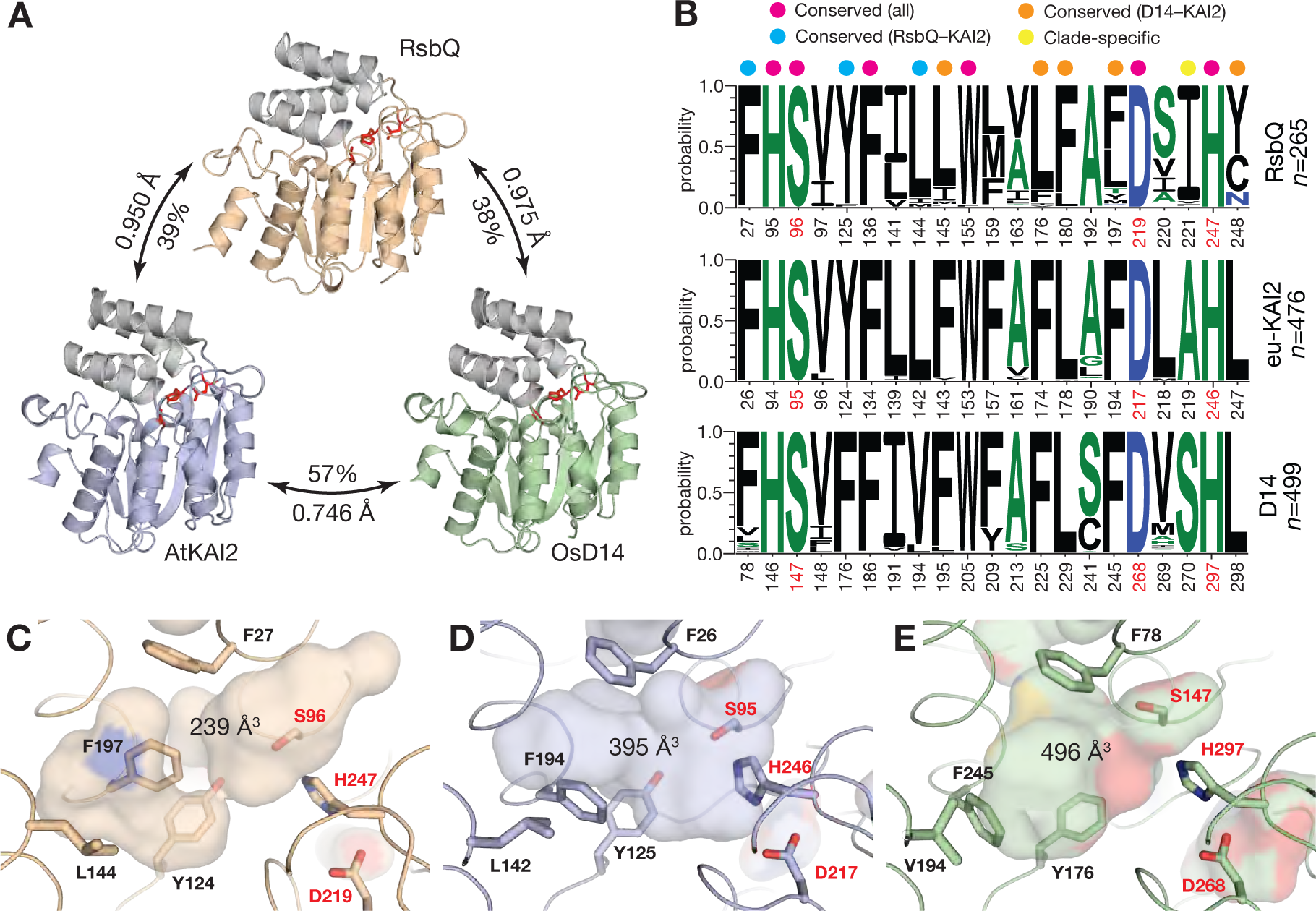
Structural comparison of bacterial RsbQ and the plant butenolide receptors KAI2 and D14. **A.** All three proteins comprise a core α/β hydrolase domain (coloured) and a lid domain of four alpha helices (grey); the catalytic triad residues are in red. Pairwise comparisons show overall percentage of amino acid identity, and RMSD values of aligned Cα atoms after outlier rejection. PDB codes: RsbQ from *Bacillus subtilis*, 1wom; AtKAI2 from *Arabidopsis thaliana*, 4hrx; OsD14 from *Oryza sativa*, 3w04. **B.** Frequency plots of amino acids at 21 positions that contribute to the ligand binding pocket among *n* RsbQ, eu-KAI2 and D14 homologues. Hydrophobic amino acids are black; neutral, green; and hydrophilic, blue. Residue numbering is per RsbQ, AtKAI2 and OsD14, with catalytic residues numbered in red. Magenta dots indicate residues that are conserved across all three groups; blue dots, conserved between RsbQ and eu-KAI2; orange dots, conserved between eu-KAI2 and D14 only; and yellow dots, specific to each group. Underlying sequence data is presented in Data S1-S4. **C–E.** Surface representation of ligand binding pockets for RsbQ (**C**), AtKAI2 (**D**) and OsD14 (**E**), highlighting catalytic residues (red labels) and selected residues (black labels) including those that are conserved between RsbQ and eu-KAI2 homologues (blue dots in B). Predicted pocket volumes are shown in cubic Angstroms.

We expressed RsbQ in *E. coli* with a 10-kDa N-terminal SUMO domain, an addition that aids solubility and is functionally benign for KAI2 and D14 proteins^6,30^. We tested the thermal stability of SUMO-RsbQ in the presence of the two constituent enantiomers of the synthetic strigolactone *rac*-GR24 (GR24^5DS^ and GR24^ent-5DS^), which contain a methylbutenolide moiety, and their desmethyl butenolide equivalents dGR24^5DS^ and dGR24^ent-5DS^ (for chemical structures, see Figure S1). KAI2 and D14 proteins are less thermally stable in the presence of GR24, a shift that likely reflects a conformational change in the lid domain that forms an interaction interface with the signalling F-box protein MAX2 ^6,7,11,30,31^. KAI2 proteins are most sensitive to desmethyl butenolides, especially dGR24^ent-5DS^, while D14 is insensitive to this compound^32,33^. In thermal shift assays, we found that SUMO-RsbQ underwent a dramatic reduction in melting temperature upon treatment with just 25 µM dGR24^5DS^, and a qualitatively similar shift with dGR24^ent-5DS^, albeit at higher ligand concentrations (Figure 2A,B). In contrast, GR24^5DS^ produced a more modest shift in melting temperature, and GR24^ent-5DS^ was inactive in this assay (Figure 2C,D). Thus, RsbQ shows a ligand preference for dGR24 over GR24 like KAI2, but with opposite stereochemical preference. To determine whether these effects are dependent on the hydrolytic function of RsbQ, we also assayed the RsbQ^S96A^ mutant. The effects of butenolides on thermal stability were much reduced or abolished entirely (Figure S2A-D), as observed with corresponding mutants of KAI2 and D14 ^10,11,33^. Furthermore, we found that SUMO-RsbQ showed greater hydrolytic activity and affinity towards desmethyl-YLG over the methylbutenolide equivalent YLG, and a greater affinity for *rac*-dGR24 over *rac*-GR24 based on intrinsic tryptophan fluorescence assays (Figure 2E,F). Although the S96A mutation greatly reduced the hydrolytic activity of RsbQ towards dYLG, it did not abolish it entirely, suggesting that this mutant retains some residual activity (Figure S2E). Notably, this mutant also retains the capacity to interact with a ligand (Figure S2A,B,F), as was reported for AtKAI2^S95A^ (ref. 33).

**Figure 2.**
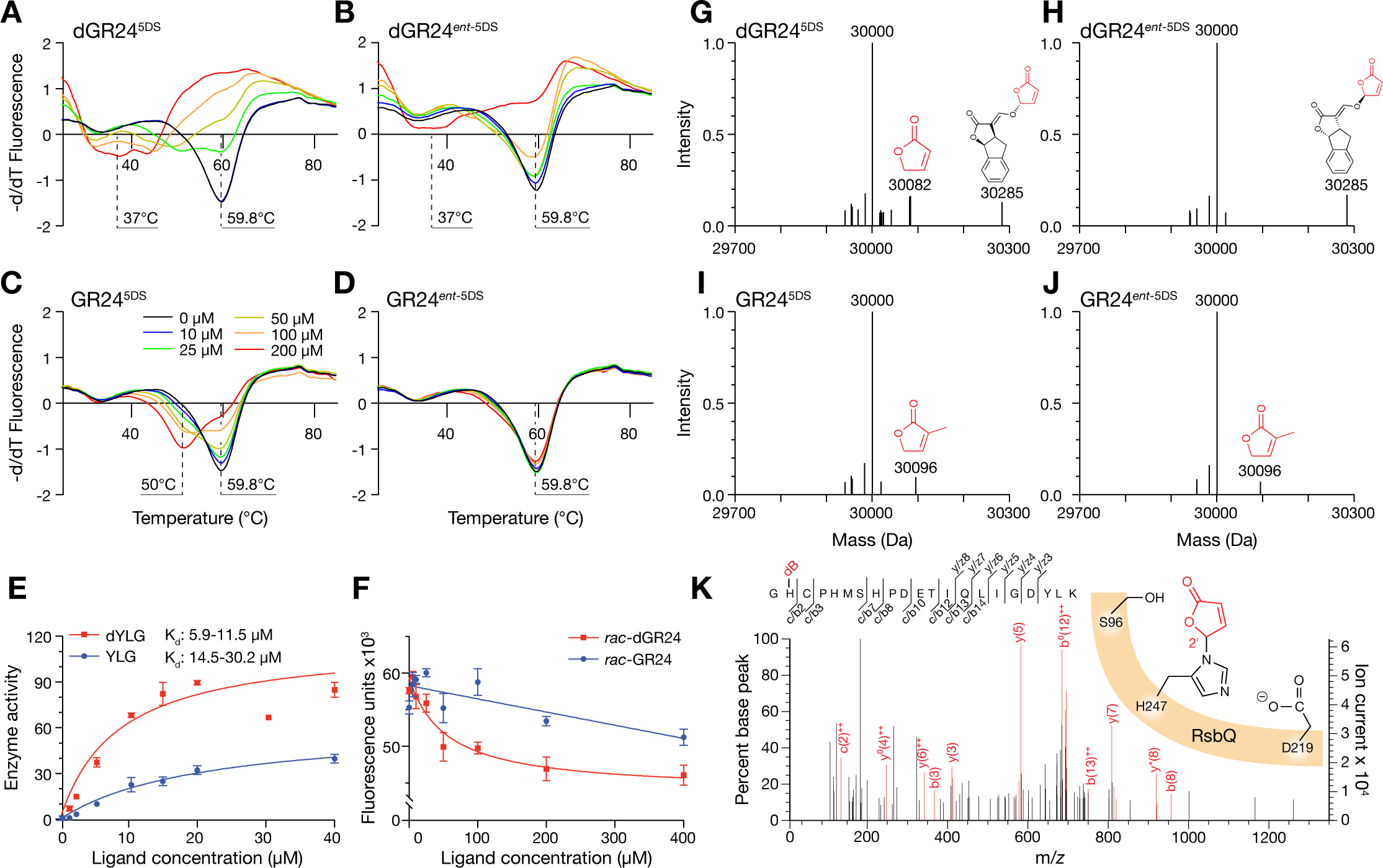
RsbQ interacts preferentially with desmethyl butenolides. **A–D.** Thermal shift assays of purified SUMO-RsbQ fusion protein challenged with increasing concentrations of the two constituent enantiomers of desmethyl-GR24 (dGR24^5DS^ and dGR24*^ent^*^-5DS^) or the two corresponding enantiomers of GR24 (C, D). Inferred protein melting temperatures are indicated with dashed lines. **E**. Hydrolysis activity of SUMO-RsbQ towards the profluorescent butenolides Yoshimulactone Green (YLG) and desme-thyl-YLG (dYLG). Data are means ± SD of n=3 technical replicates. K_d_ values were estimated by non-linear regression and are given as 95% confidence intervals. **F**. Intrinsic tryptophan fluorescence measurements of SUMO-RsbQ in response to varying concentrations of racemic GR24 or dGR24. Data are means ± SD of n=4 technical replicates. **G–I**. Deconvoluted intact mass spectra of purified native RsbQ (calculated MW=30,000 Da) treated with individual enantiomers of dGR24 and GR24. Shifted peaks are labelled with observed MW and the chemical structure of putative adducts consistent with the treatment and increase in mass. **K**. Tandem mass spectrum of a fragmented RsbQ peptide (Gly246 to Lys266; M_mi_ = 2390.1 Da) following incubation with *rac*-dGR24 and trypsin digestion. A peptide (M_mi_ = 2472.1 Da) was identified using MASCOT (version 2.5.1, Matrix Sciences) with ions (red peaks) consistent with a RsbQ peptide carrying a desmethyl butenolide ring (dB, (+82.0)) on the catalytic His247. Identified peaks representing c/b and y/z ions are annotated with ^++^, indicating a doubly-charged ion; ^0^ representing an ion − H_2_O and * an ion − NH_3_. Cartoon indicates the putative dB modification (red) on the His247 residue.

Butenolide signalling via KAI2 and D14 involves ligand hydrolysis, which initiates with nucleophilic attack upon the butenolide carbonyl group by the catalytic serine. This mechanism is supported by the experimental detection of covalently modified intermediates, including a histidine adduct consistent with the butenolide group^4,5,34^. Using intact denatured mass spectrometry of purified, native RsbQ treated with dGR24^5DS^, we observed a peak corresponding to a mass increase of +82 Da, which is equivalent to an adduct derived from the desmethyl butenolide moiety (Figure 2G). We did not observe this peak with dGR24^ent-5DS^, but saw +96 Da peaks with both enantiomers of GR24 that were consistent with a methyl-substituted butenolide adduct (Figure 2H–J), as reported for D14 and KAI2 homologues^4,5^. Unexpectedly, we also observed a peak of +285 Da when treating with both enantiomers of dGR24, a mass that closely corresponds to an intact dGR24 molecule. This peak was also observed with RsbQ^S96A^, while the +82/96 Da peaks were absent (Figure S2I–L). Thus, the formation of the +82/96 peak requires the catalytic serine and likely reflects an intermediate of ligand hydrolysis, but the +285 Da peak is a non-specific adduct of uncertain location that is unique to dGR24. Tandem mass spectrometry of trypsin-digested RsbQ treated with *rac*-dGR24 identified a peptide consistent with a covalent modification of the catalytic histidine residue with a desmethyl butenolide moiety of mass 82 Da (Figure 2K; Table S1). Notably, we did not observe a similar modification of RsbQ mock-treated with DMSO (Figure S2M, Table S1). Overall, these results indicate that RsbQ has a preference for desmethyl butenolide ligands and operates with the same catalytic mechanism as KAI2 and D14 from plants.

We sought to characterise the response of *B. subtilis* to butenolides that are bioactive in plants. The *amyE::ctc-LacZ* transgene drives β-galactosidase expression from the σ^B^-dependent *ctc* promoter^35^. Consistent with single-cell RNA sequencing^36^, we observed that σ^B^ activity peaked at the exit from exponential growth (OD_600_ ∼1.0) and was sustained throughout the transition to stationary phase; notably, this peak was absent in the null *ΔrsbQ* mutant (Figure S3A). Unexpectedly, we observed that *rac*-dGR24 and *rac*-GR24 strongly inhibited the early induction of σ^B^, with *rac*-dGR24 being at least 10-fold more potent (Figure 3A). The effects of GR24 and dGR24 were not fully sustained, as β-galactosidase activity partially recovered at later phases of growth (Figure S3B), which may have resulted from metabolism and/or chemical hydrolysis of the compounds over time. The desmethyl debranone *rac*-dCN-deb was 10-fold weaker even than GR24, while the methylbutenolide equivalent *rac*-CN-deb was inactive (Figure S3C); this pattern was also reflected in thermal shift assays (Figure S3E,F). Karrikins also had no effect on β-galactosidase activity (Figure S3D), consistent with observations that karrikins are inactive *ex planta*^37–40^. The inhibitory effect of GR24 and dGR24 on the *amyE::ctc-LacZ* reporter was transcriptional (Figure 3B), suggesting that these inhibitory effects resulted from a general decrease in σ^B^ activity, and not from an indirect effect on β-galactosidase.

**Figure 3.**
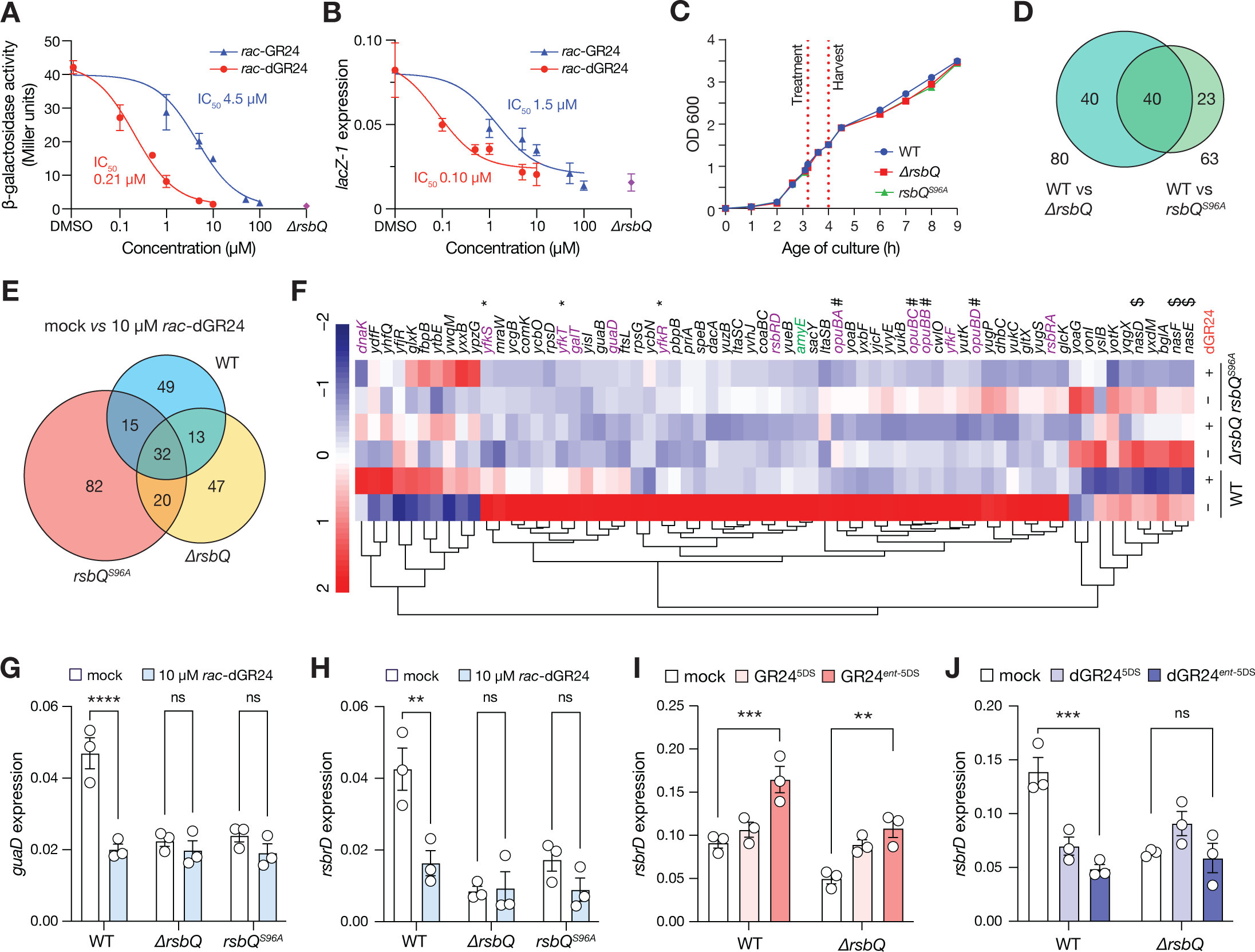
*Bacillus subtilis* responds to desmethyl butenolides through an RsbQ-dependent mechanism. **A-B**. Response of β-galactosidase reporter activity (A) and *lacZ-1* transcripts (B) in *B. subtilis* 168 carrying the *amy-E::ctc-LacZ* transgene to treatment with varying concentrations of racemic GR24 and dGR24. Corresponding values of the untreated *ΔrsbQ* mutant are shown in violet. Cultures were treated upon reaching OD_600_ of 0.5, and harvested 1 hour later (OD_600_ ∼1.2). Data are means ± SE n=3 replicate cultures grown at 37 °C on separate days; data in A and B are derived from the same cultures. *lacZ-1* transcripts were normalised to *gyrB* and *recA* reference transcripts. IC_50_ is the inferred concentration for 50% inhibition. **C.** Growth curves of WT and mutant strains in LB medium under conditions for RNA-seq (37 °C 180 rpm). Dashed lines indicate treatment and harvest based on time after inoculation. Data are means ± SE of three replicate cultures. **D.** Overlap of differentially expressed genes (DEGs) in cultures mock-treated with 0.1% DMSO, comparing WT with each of the two *rsbQ* mutants. DEGs were defined by log_2_ fold change >1 or <−1, and false discovery rate-adjusted P-value < 0.05. Data are derived from three replicate cultures per genotype/treatment condition. **E.** Overlap of DEGs in each genotype treated with 0.1% DMSO (mock) versus 10 µM *rac*-dGR24 (using the same selection criteria as **D**). For example, 49 genes were differentially expressed in WT only following *rac*-dGR24 treatment. **F.** Hierarchical clustering analysis of 47 RsbQ-dependent DEGs and 20 *rac*-dGR24-specific DEGs (using the same selection criteria as **D**). Blue and red represent downregulation and upregulation, respectively. Z-scores were calculated from normalised count data. Genes in purple are known members of the σ^B^ regulon. Genes with the same symbol belong to the same operon. The *ctc-LacZ* reporter present in all three genotypes is integrated into the *amyE* locus; spurious differential expression of *amyE* (highlighted green) reflects readthrough of the reporter, and is not counted in the set of 67. **G–H**. Levels of RsbQ-dependent *guaD* and *rsbrD* transcripts in response to 10 µM *rac*-dGR24 as determined by qRT-PCR from an experiment independent from the RNA-seq dataset. **I–J**. Levels of *rsbrD* transcripts following treatment with the constituent enantiomers of GR24 (**I**) or dGR24 (**J**) at 5 µM. In **G–J**, cultures were treated and harvested as shown in **C**; transcripts were normalised to *gyrB* and *recA* reference transcripts. Data are means ± SE of n=3 replicate cultures. Significant differences are indicated: ** P<0.01; *** P<0.001; **** P<0.0001; ns, non-significant (two-way ANOVA).

To understand in more detail the effects of butenolides upon RsbQ cellular function, we undertook a transcriptome analysis of WT, *ΔrsbQ* and *rsbQ^S96A^* in response to treatment with 10 µM *rac*-dGR24. Treatment was timed to coincide with peak σ^B^ activity at late exponential growth (Figure 3C). Initially we compared WT with each mutant without *rac*-dGR24 treatment and found a total of 103 differentially expressed genes across both comparisons; of these, only 40 (38%) were common (Figure 3D; Table S2). The *ΔrsbQ* and *rsbQ^S96A^* mutant transcriptomes were relatively similar under mock-treatment conditions, but diverged in their transcriptomic response to *rac*-dGR24 (Figure 3E, Figure S4A,B; Table S3). This finding implies that the two mutant alleles are phenotypically distinct, with *ΔrsbQ* being the stronger of the two, and echoes the residual activity of the RsbQ^S96A^ protein we observed *in vitro*. Regardless, we defined these shared 40 genes as RsbQ-dependent, while recognising that this may be an under-representation.

A total of 109 genes were differentially expressed in WT following treatment with *rac*-dGR24 (Figure 3E, Figure S4C). Of these, only 49 (45%) were unique to WT and therefore dependent on RsbQ (Table S3). Overlaps between this set and the 40-gene RsbQ-dependent set meant that a total of 67 unique “core RsbQ genes” were robustly responsive to *rac*-dGR24 in an RsbQ-dependent manner (Table S4). Hierarchical clustering of these genes across the six treatment/genotype combinations showed that treatment of *B. subtilis* with *rac*-dGR24 phenocopies loss of RsbQ (Figure 3F), which is fully consistent with the inhibitory effect of butenolides on the *amyE::ctc-LacZ* reporter and the positive effect of RsbQ on σ^B^ activity. Of these 67 genes, 13 (19%) are members of the σ^B^ regulon; given that this regulon comprises ∼255 of the 4174 (6%) protein coding genes in *B. subtilis*^21^, this represents a significant enrichment (P=0.0001, binomial test, two-tailed). Nevertheless, the small subset of the σ^B^-regulon that is under direct control of RsbQ likely reflects the substantial diversity in promoter motifs and expression profiles among σ^B^ targets, which are regulated by multiple inputs^21^.

We confirmed the inhibitory effects of *rac*-dGR24 on transcript levels of several of the 13 σ^B^- and RsbQ-dependent genes, in a separate experiment, using qRT-PCR (Figure 3G,H; Figure S5). We then compared the effects of individual enantiomers of GR24 and dGR24 on these genes. Surprisingly, we found that neither GR24 enantiomer could repress the transcripts that we assayed, and instead moderately promoted their expression in an RsbQ-independent manner (Figure 3I, Figure S6). Thus, even if GR24 does influence RsbQ signalling, it also appears to have independent and confounding effects. In contrast, the dGR24 enantiomers affected levels of RsbQ-dependent transcripts in WT only, consistent with the RNA-seq filtering criteria. Both dGR24^5DS^ and dGR24^ent-5DS^ repressed the expression of genes such as *rsbrD*, *guaA* and *yfkS*, but dGR24^ent-5DS^ was typically the more active enantiomer (Figure 3J, Figure S6). This result appears to contradict the relative activities of the two enantiomers *in vitro* (Figure 2), although discrepancies between bioactivity and *in vitro* activities have been reported for KAI2 and D14 proteins. For example, dGR24^5DS^ exhibits no bioactivity in *Arabidopsis thaliana*, but nevertheless is hydrolysed efficiently by both AtKAI2 and AtD14, and induces thermal instability of purified AtKAI2^33^. Likewise, the *S*-configured enantiomer of CN-deb is bioactive via AtKAI2, but does not perceptibly affect its thermal stability^33,41^.

The nutritional stress pathway mediated by RsbQ and RsbP negatively regulates biofilm formation via σ^B^ in suitable strains of *B. subtilis* ^20,42^. To determine if butenolides can interfere with this process, we assayed biofilm formation in stationary cultures of the non-domesticated strain NCIB3610 *comI^Q12L^* treated with varying concentrations of *rac*-GR24 and *rac*-dGR24. We found that biofilm production was enhanced by the addition of both compounds and in a dose-dependent manner: at higher concentrations, biofilm production approached that produced by a *ΔrsbQP* knockout mutant (Figure 4). Statistically significant increases in biofilm production were triggered by 5 µM *rac*-dGR24, compared with 50 µM for *rac*-GR24, representing ∼10-fold higher potency in this assay. These results suggest that, at least in *B. subtilis*, exogenous butenolides can influence the transition from free-living growth to a community lifestyle adapted to growth on surfaces.

**Figure 4.**
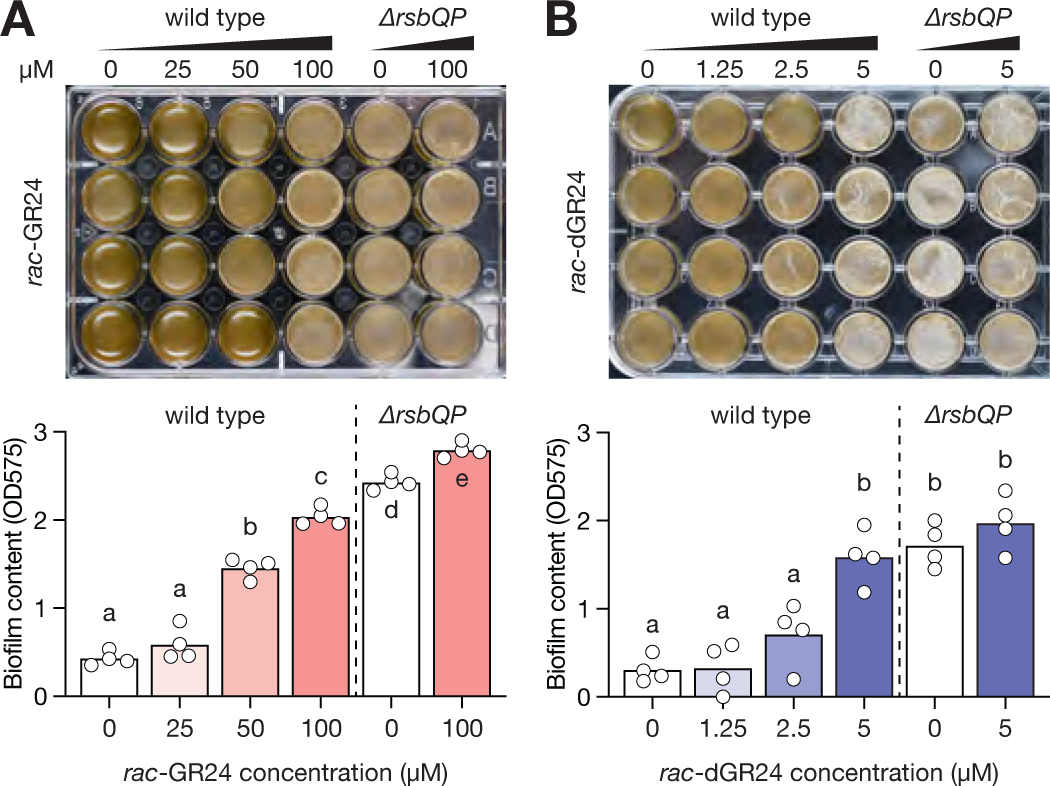
Butenolides promote biofilm formation in NCIB3610 via RsbQP. Biofilm formation in static cultures of the undomesticated *B. subtilis* strain NCIB3610 *ComI^Q12L^* and a *ΔrsbQP* deletion mutant of the same strain. Upper panels show representative 24-well assay plates after 22-24 h of growth at 30 °C in the dark. Lower panels show quantification of biofilm content using crystal violet stain. Biofilm levels were assessed in response to varying concentrations of *rac*-GR24 (**A**) and *rac*-dGR24 (**B**). Bars are means; individual data points are derived from individual wells of the assay plate (n = 4 wells per treatment). Different lower-case letters indicate a significant difference between treatments (one-way ANOVA, P<0.05). This experiment was repeated at least twice with similar results.

Our results reveal that RsbQ, the *B. subtilis* homologue of the plant karrikin and strigolactone receptors, mediates responses to butenolide compounds that are bioactive in plants. These findings suggest the exciting possibility of interkingdom communication mediated by butenolide signals. Besides *Bacillus* sp., the RsbQ-P module is prevalent among diverse bacteria associated with plants, including *Rhizobium leguminosarum*, *Mesorhizobium* sp., *Pseudomonas syringae* and *Paraburkholderia phytofirmans*^9^. Strigolactones exuded by pea roots promote nodulation with *R. leguminosarum*, most probably by acting on the bacterial partner ^43^. However, a recent study on pea but with a different bacterial strain reached a contradictory conclusion in which strigolactones appeared to repress nodule development and maturation^44^. There is also potential for butenolide signals derived from bacteria: for example, *Streptomyces albus* J10764 secretes a series of desmethyl butenolides that can stimulate interspecific secondary metabolism in *S. avermitilis*^45^, and at least one of these compounds can promote seed germination of the parasitic weed *Orobanche minor*^46^. An ancestral role for strigolactones as rhizosphere signals to recruit arbuscular mycorrhizal symbionts is likely^47^, supporting an ancient function for butenolides as a means of communication among plants and microbes. Consistent with this view, a strigolactone-deficient mutant of rice has a distinct rhizomicrobiome that is depleted in Acidobacteria, which are generally considered to have plant growth-promoting properties^48,49^. We found a clear link between biofilm growth and the inhibition of RsbQ-dependent signalling by butenolides. Although this effect may be specific to *B. subtilis*, this result reveals a direct route by which plant metabolites may promote bacterial colonisation of roots, and thereby reinforce specific bacterial composition in the rhizosphere. A wider examination of loss of RsbQ-mediated signalling in plant-associated bacteria is warranted.

It is especially noteworthy that exogenous butnolides inhibited RsbQ *in vivo*. Because RsbQ is a positive regulator of σ^B^, we assume that any endogenous ligand of RsbQ is an activator, rather than an inhibitor. Thus, dGR24 and GR24 cannot be true mimics of an endogenous RsbQ ligand, instead behaving as competitive inhibitors that “poison” RsbQ and block its subsequent function. The conformational change in RsbQ induced by dGR24 – inferred by thermal shift assays – is presumably unsuitable for promoting a protein-protein interaction with RsbP, which is arguably now the most plausible model for how the energy stress signal is transduced^23,27^. There is no obvious route for the biosynthesis of an endogenous RsbQ ligand, as none of the *Rsb* loci encode plausible biosynthetic enzymes. Perhaps without coincidence, no biosynthetic pathway for KL in plants has been found by genetic approaches. Desmethyl butenolides are the most potent compounds for inhibiting RsbQ function in *B. subtilis*, which is consistent with the endogenous RsbQ ligand and KL having a similar structure and metabolic origin. As such, it should be considered that both compounds are produced non-enzymically, possibly from common cellular metabolites.

## Supporting information

supp_tables_all

## ACKNOWLEDGEMENTS

We thank Chet Price (UC Davis) for provision of *rsbQ* mutant strains; Adrian Scaffidi (UWA) for synthesis and separation of butenolide chemicals; and Adil Khan (Texas Tech University) for guidance on transcriptome analysis. Funding was provided by the Australian Research Council to MTW, GF and GRF (DP210103078). KTM, JY and MH were supported by Research Training Program scholarships from the Australian Government. MC was supported by a Future Science Platform SynBio Fellowship from CSIRO. The Australian Genomics Research Facility provided RNA sequencing services. Mass spectrometry was performed on instruments managed by the Western Australian Proteomics Facility (WA Proteomics and Proteomics International) as a node of Proteomics Australia and was supported by infrastructure funding from the Western Australian State Government in partnership with Bioplatforms Australia under the Commonwealth Government National Collaborative Research Infrastructure Strategy.

## AUTHOR CONTRIBUTIONS

Conceptualization, MTW; Methodology, KTM, MK, MTW; Investigation, KTM, MK, JY, MH, MC, NLT, MTW; Resources, NLT, MC, GF and GRF; Writing – Original Draft, KTM, MK, MTW; Review and Editing, MTW; Visualisation, KTM, MK, MTW; Funding Acquisition, MTW, GF and GRF; Supervision, MTW, GF, and GRF.

## RESOURCE AVAILABILITY

Further information and requests for resources and reagents should be directed to the lead contact, Mark Waters.

## DATA AND CODE AVAILABILITY

The RNA-seq data are deposited in the GEO database under the reference (GSE236759). This paper does not report original code. Any additional information required to reanalyse the data reported in this paper is available from the lead contact upon request.

## EXPERIMENTAL MODEL AND SUBJECT DETAILS

### Strains, media, culture, and treatment conditions

Bacterial strains are listed in Table S5. Unless specified otherwise, strains were grown in LB (1% [w/v] tryptone, 0.5% [w/v] yeast extract and 1% [w/v] NaCl). For all experiments, the initial inoculum was from a fresh colony struck from a −80°C freezer stock onto LB 1.5% [w/v] agar plates and incubated overnight at 37 °C.

For purification of recombinant protein, expression plasmids were transformed into Rosetta and selected on LB-agar plates containing 100 µg/mL carbenicillin and 34 µg/mL chloramphenicol. Three colonies from each plate were inoculated into a single vial containing 5 mL LB supplemented with the same antibiotics and incubated overnight at 37 °C/180 RPM to generate a starter culture. The starter culture was diluted 1:1000 into 450 mL LB in a 2-L Erlenmeyer flask, supplemented with 100 µg/mL carbenicillin, and grown at 30 °C/180 RPM until OD_600_ ∼0.75-1.0, at which point 0.1 mM isopropyl β-D-1 thiogalactopyranoside was added to induce protein expression. The culture was incubated at 16 °C/180 RPM for a further 17 hours. Cells were collected by centrifugation at 5,000×*g* for 15 minutes, and cell pellets were stored at −80 °C.

For the β-galactosidase reporter assay, *B. subtilis* strains PB198 and PB605 carrying the *amyE::ctc-LacZ* reporter construct were cultured on LB plates containing 7 µg/mL chloramphenicol. Three colonies were used to inoculate a 5-mL culture with antibiotics, grown overnight at 30 °C/180 RPM. The starter culture was then diluted 1:1000 into 50 mL LB in a 250 mL Erlenmeyer flask with antibiotics, and incubated at 37 °C/200 RPM. Butenolide treatments began at OD_600_ 0.5 by adding 125 µL of 400× concentrated stock dissolved in DMSO. For time course experiments, 500-µL samples were taken in triplicate every 30 minutes, centrifuged at 15,000×*g* for one minute, washed with 500 µL of 50 mM TRIS-HCl pH 7.5 and re-pelleted with a second centrifugation. Cell pellets were frozen in liquid nitrogen and stored at −80 °C. For dose-response experiments with *rac*-dGR24 and *rac*-GR24, samples were treated at OD_600_ 0.5 and grown for 1 hour before collecting samples. For dose-response experiments with *rac*-debranones and karrikins, samples were grown at 30 °C/200 RPM, treated at OD_600_ 0.5 and grown for 2 hours before collecting samples.

For qRT-PCR and RNA-seq studies, strains PB198, PB605 and PB634 were cultured on selective plates containing 7 µg/mL chloramphenicol. Three colonies were used to start a 5-mL culture with antibiotics, grown overnight at 37 °C/185 RPM. To initiate the experiment, approximately 500 µL (1:100) of the starter culture was inoculated into 50 mL of prewarmed liquid LB in a 250 mL Erlenmeyer flask and grown in a shaker incubator (Innova 42, New Brunswick Scientific) at 185 RPM and 37 °C. Optical density (OD_600_) was monitored throughout the experiment. Pilot experiments based on *ctc* transcript levels were used to determine optimal times for treatment and harvest. Cultures were treated with either DMSO (mock) or 10 µM *rac*-dGR24 at OD_600_ 1.0–1.1 (Figure 3C). Once cultures achieved OD_600_ 1.4–1.5, they were promptly centrifuged at 10,000×*g* for 2 minutes at 4 °C. Pellets were frozen in liquid nitrogen. For RNA-seq, three independent experimental replicates were collected on separate days. Each replicate consisted of six samples comprising the three genotypes (PB198, PB605 and PB634) and both treatments (mock, 10 µM *rac*-dGR24).

For biofilm assays, *B. subtilis* strains DK1042 and WLB042 were streaked from glycerol stocks on selective plates (no antibiotics for DK1042 and 7 µg/mL chloramphenicol for WLB042). A non-biofilm-producing colony was used to inoculate a 5 mL LB starter culture, grown at 30 °C/180 RPM for 16 hours. The following day, 40 mL LBB (LB supplemented with 4% yeast extracts) media in a 50 mL centrifuge tube was inoculated with 1:250 dilution of starter culture (without antibiotics). Four replicate aliquots, each of 2 mL, were then transferred to a 24-well polystyrene plate (Sarstedt 83.3922). Each well was treated with 5 µL ligand at the required concentration (from 400× stocks dissolved in DMSO). For growth, the plate was placed in a sterilised plastic food container with ventilation holes drilled in the lid, and sealed with breathable tape (Leukopor). The setup was then incubated at 30 °C under static and dark conditions for 22–24 hours when biofilm development was observed. Photographs of biofilm were taken at the end of the experiment.

## METHOD DETAILS

### Sequence conservation analysis

RsbQ homologues were identified by TBLASTN searches against the NIH nr/nt database using *B. subtilis* RsbQ (UNIPROT O07015) as a query, limiting search set to Bacillota (taxid: 1239) and excluding Bacillus subtilis group (taxid: 653685). Expect threshold was set to 0.01, and results were filtered to percent identity >40%, E-value <1e-50. KAI2 and D14 homologues were identified by TBLASTN searches against the 1000 Plant Transcriptomes ‘onekp v5’ database (https://db.cngb.org/onekp/) using *Arabidopsis thaliana* KAI2 (UNIPROT Q9SZU7) and D14 (UNIPROT Q9SQR3) as queries. Parameters were maximum 500 target sequences; expect value <0.01; word size of six; and BLOSUM62 scoring matrix. The KAI2 dataset was the same as that described previously^30^. Nucleotide sequences were imported into Geneious R10 software and searched for ORFs. Truncated sequences and duplicate ORFs were removed. Protein sequences were aligned using the MAFFT algorithm (BLOSUM62, all values default). Clearly divergent or frame-shifted sequences were removed. The alignment was then trimmed to leave the 21 selected pocket residues, and this alignment was uploaded to WebLogo3 (https://weblogo.threeplusone.com/create.cgi). A list of analysed sequences and their amino acid alignments are provided in Data S1-S4.

### Molecular cloning

Construction of pSUMO-RsbQ/RsbQ^S96A^: The coding sequences for RsbQ and RsbQ^S96A^ were PCR amplified from strains PB198 and PB634 respectively, transferred into the pE-SUMO-Amp vector by Hot Phusion assembly^50^, and verified by Sanger sequencing.

Construction of pMDY001c: Homology arms comprising 1000 bp upstream and 1000 bp downstream of the *rsbQP* operon with an intervening spectinomycin resistance cassette were PCR-amplified and inserted by Hot Fusion assembly into pBS4S, replacing the existing homology arms for integration into the *thrC* locus. Subsequently, the Spec^R^ resistance cassette was replaced with a Cm^R^ resistance cassette isolated from pBS3C-lux using restriction-ligation. Plasmids are listed in Table S5. Cloning primers are listed in Table S6.

### Transformation of *B. subtilis*

Freshly-streaked colonies were resuspended in 5 mL of Medium A (15.1 mM (NH4)_2_SO_4_, 80 mM K_2_HPO_4_, 44 mM KH_2_PO_4_, 3.4 mM Na_3_C_6_H_5_O_7_, 0.45% (w/v) glucose, 0.019% (w/v) tryptophan, 5.74 mM MgSO_4_, 0.11% (w/v) casamino acids and 1 µg/mL (NH_4_)_5_[Fe(C_6_H_4_O_7_)_2_]), and grown at 37 °C/180 RPM until optical density reached an OD_600_ of 0.2. Growth then continued for three additional hours. Following this growth period, an equal volume of Medium B (15.1 mM (NH_4_)_2_SO_4_, 80 mM K_2_HPO_4_, 44 mM KH_2_PO_4_, 3.4 mM Na_3_C_6_H_5_O_7_, 0.45% (w/v) glucose and 5.74 mM MgSO_4_) was added, and the culture was incubated for two further hours. Concurrently, plasmids were linearised by digestion with *Sca*I. For the transformation, 400 µL of the *B. subtilis* culture was mixed with 500 ng linearised DNA in a 2-mL microfuge tube, followed by a 1 h incubation at 37 °C/180 RPM. The mixture was plated on selective LB-agar plates. Transformed colonies were verified for successful recombination by PCR-amplification of both the upstream and downstream connections between the insert and bacterial genome. Primers are listed in Table S6.

### Protein purification

Cell pellets were thawed, resuspended in 25 mL of lysis buffer (20 mM HEPES, 150 mM NaCl, 10% (v/v) glycerol, 40 mM imidazole, 1× BugBuster reagent [Novagen], and 25 U/mL Benzonase nuclease [Novagen], pH 7.5), and incubated at room temperature on a reciprocal shaker for 15 minutes. The suspension was clarified by centrifuging twice at 15,000×*g* for 20 minutes. The clear lysate was then loaded on a 30 mL Econo-Pac gravity chromatography column (Bio-Rad) containing 2 mL settled Ni-NTA resin (Bio-Rad) pre-equilibrated with wash buffer (20 mM HEPES, 150 mM NaCl, 10% (v/v) glycerol, 40 mM imidazole, pH 7.5). The column was incubated with lysate at 4 °C with rotation for one hour, after which the column was allowed to drain under gravity. The column was washed twice with 20 mL wash buffer at 4 °C, and the bound protein was eluted with 8 mL of elution buffer (20 mM HEPES, 150 mM NaCl, 10% (v/v) glycerol, 200 mM imidazole, pH 7.5). For removal of the 6×HIS-SUMO tag, the eluted protein was digested overnight with the SUMO-protease UlpI (ref 51) (1 µg UlpI per 200 µg SUMO-RsbQ) and simultaneously dialysed against 30-fold volume of wash buffer for 16 hours at 4 °C. The following day, the samples were passed through a gravity chromatography column containing 2 mL Ni-NTA pre-equilibrated with wash buffer and incubated for one hour at 4 °C with rotation. The flow-through, containing the tag-removed protein, was collected. Imidazole was removed via repeated concentration and dilution steps with 20 mM HEPES, 150 mM NaCl, 10% (v/v) glycerol using an Amicon Ultra-4 centrifugal column with a 10 kDa molecular weight cut-off (Merck-Millipore). This process yielded a protein concentrated to ∼10–20 mg/mL. Protein concentration was estimated by A_280_, and purity assessed by SDS-PAGE.

### Intact denatured protein mass spectrometry

A 25-µL reaction containing 50 µM purified RsbQ in 20 mM HEPES, 150 mM NaCl (pH 7.5) was treated with 0.625 µL of 40× concentrated ligand in DMSO (final conc. 50 µM dGR24, or 500 µM GR24). dGR24-treated samples were incubated for 10 minutes at room temperature, while GR24 treated samples were incubated for 30 minutes at room temperature. Reactions were quenched with seven volumes (175 µL) of MS buffer (2% [v/v] acetonitrile (MeCN), 0.1% [v/v] formic acid (FA) in MS-grade water). Samples were analysed on a ThermoScientific Q Exactive HF Orbitrap LC-MS/MS System (ESI in positive ion mode with an in-source CID of 25 eV, a resolution of 15000 and a scan range of 400 to 4000 m/z, nebulizer gas (N_2_), 1.4 bar; dry gas (N_2_), 8 L/min and dry temperature, 200 °C) connected to Waters XBridge Protein BEH C4 Column (300 Å particle size; 3.5 µm pore size; 2.1 mm × 100 mm; column oven 60 °C). UHPLC elution used a flow rate of 0.15 mL/min with buffers A (2% [v/v] MeCN, 0.1% [v/v] FA) and B (98% [v/v] MeCN, 0.1% [v/v] FA). The elution profile was: 0-5 min, 95% A & 5% B; 5-35 min, a linear gradient to 2% A & 98% B; 35-40 min, 2% A & 98% B; and 40-45 min, a linear gradient back to 95% A & 5% B. Acquired spectra were analysed and deconvoluted using UniDeck software as described previously^52^, using UniChrom2 on isotopic resolution pre-set with modified settings for Picking range (1 Da) and Picking threshold (0.07).

### Digested protein mass spectrometry

Purified RsbQ (7 µL at 467 µM in 20 mM HEPES, 150 mM NaCl, 10% (v/v) glycerol, pH 7.5) was treated with 3 µL of 3.26 mM *rac*-dGR24 in 10% [v/v] DMSO, achieving a 1:3 protein:substrate ratio. Reactions were left for 15 minutes at room temperature. To quench the reaction, four volumes of pre-chilled acetone (–20 °C) were added, followed by a two-hour incubation at –20 °C. The precipitated protein was then centrifuged at 20,000×*g* for 20 minutes at 4 °C, the supernatant was pipetted off, and residual acetone further evaporated in a vacuum centrifuge for 10 minutes. The sample was resuspended in 20 µL STT buffer (4 % SDS (w/v), 0.5 M Tris(2-carboxyethyl)phosphine [Bond-Breaker™ Solution Neutral pH, Pierce], 0.1 M Tris-HCL, pH 7.6) and incubated at 95 °C for 30 minutes. Procedures for sample clean-up and trypsin digestion were followed according to Mikulášek et al. 2021 (ref 53). Samples were treated with 20 mM iodoacetamide and incubated in the dark at ambient temperature for 30 minutes, followed by 5 mM DTT for 10 minutes, to eliminate cysteine disulfide bonds. A 5-µg aliquot of protein was bound to SP3 beads and mixed with 50% (v/v) ethanol at 1000 RPM for 10 minutes on a heated shaking incubator (Eppendorf ThermoMixer C) at room temperature. The supernatant was removed, and the beads were washed five times with 140 µL 80% (v/v) ethanol, with transfer to a fresh tube in the last wash. Sample digestion was carried out by resuspending beads in 1 µL of trypsin solution (0.2 µg trypsin in Promega V5111 resuspension buffer) and 19 µL 50 mM ammonium bicarbonate and incubating at 37 °C with agitation for 18 hours. The digestion was quenched with 1 µL of 10% (v/v) formic acid, and the sample centrifuged twice at 20,000×*g* for 10 minutes, removing SP3 beads by transferring the supernatant to a new tube each time.

For mass spectrometry analysis ∼0.1 µg protein was injected into an online nanoflow (200 nL min^-1^) 500 mm capillary column (Picofrit with 10 μm tip opening and 75 μm diameter; New Objective, PF360–75-10-N-5) packed in-house with 150 mm of C18 silica material (3 µm; Dr. Maisch GmbH). The column was connected to a ThermoScientific Orbitrap Exploris 480, inline with Dionex Ultimate 3000 series UHPLC. Spray voltage was set to 1.9 kV, and heated capillary temperature at 250 °C. Mobile phase buffer A was composed of 0.1% [v/v] FA. Mobile phase B was composed of 100% [v/v] ACN, 0.1% [v/v] FA. Each sample was separated over a 33-minute gradient, including time for column re-equilibration. Flow rates were set at 200 nL/min. Full MS resolution was set to 60,000 at m/z 200 and full MS automatic gain control (AGC) target was set to standard with an injection time of 25 ms. Mass range was set to 350–1400. AGC target value for fragment spectra was set at 100% with a resolution of 15,000 and injection times of 28 ms. Intensity threshold was set at 2X10^4^, isolation window at 1.4 m/z, and normalized collision energy at 30%.

Data analysis was carried out using Mascot (Matrix Science). The default Thermo .raw files were converted to .mzml format using msconvert software (https://bioinformaticshome.com/tools/proteomics/descriptions/msconvert.html#gsc.tab=0) and searched in Mascot against a *Bacillus subtilis* (Strain 168) database comprising the Uniprot Taxon 224308 proteome containing a total of 8538 protein sequences. Searches were conducted using the Mascot search engine version 2.1.04 (Matrix Science) utilizing error tolerances of ± 100 ppm for MS and ± 0.5 ppm for MS/MS, ‘Max Missed Cleavages’ set to 1, and Oxidation (M), Carboxymethyl (C) as variable modifications. The Instrument was set to ESI-Trap and Peptide charges set at 2+, 3+ and 4+. Results were filtered using ‘Standard scoring’, ‘Max. number of hits’ was set to 20, ‘Significance threshold’ at p< 0.05 and ‘Ions score cut-off’ at 0. Searches examining for desmethyl-butenolide modifications included a variable monoisotopic mass of 82.005479.

### Thermal shift assays

Protein and ligand solutions were prepared separately. Protein solution comprised of 40 µM protein in a buffer containing 20 mM HEPES, 150 mM NaCl and 10× SYPRO tangerine reporter dye (diluted 1:500 from the commercial 5000× stock in DMSO), pH 7.5. The ligand solution was prepared at double the desired concentration by diluting it 1:10 from a 20× stock solutions in acetone, using a buffer of 20 mM HEPES, 150 mM NaCl pH 7.5. To set up the assay, 5 µL of each solution was combined in a 384-well microtiter plate, resulting in a 10 µL reaction mix comprising 20 µM protein, 5× SYPRO Tangerine dye, 20 mM HEPES, 150 mM NaCl, 5% (v/v) acetone and 0–200 µM ligand. The plate was sealed, centrifuged at 1000×*g* for two minutes at 4 °C and subsequently dark-incubated for 30 minutes at ambient temperature. Analysis was done on Roche LC480 thermocycler with the temperature gradually increasing from 20 °C to 80 °C at a rate of 0.2 °C/s. Measurements were taken by using excitation at 483 nm and emission of 640 nm at a rate of 20 acquisitions/min. Data was retrieved from LC480 software using the “Tm calling” function plotting temperature against the derivative of 640 nm fluorescent measured over temperature for each sample. Data from four replicate samples were imported to GraphPad Prism and averaged to generate the final graphs.

### YLG/dYLG hydrolysis assays

Purified SUMO-RsbQ was diluted to 0.1 µg/µL in reaction buffer (20 mM HEPES, 150 mM NaCl, pH 7.5). Stocks of Yoshimulactone Green (YLG) or desmethyl-YLG (dYLG) dissolved in DMSO were serially diluted to 1,000× final concentration. These were subsequently diluted in reaction buffer to yield a 1.11× final concentration. Each sample was prepared in a black 96 well microtiter plate (Greiner 655900). Using a repeat pipettor, 10 µL of protein solution, containing 1 µg SUMO-RsbQ, was dispensed into each well. Using a multichannel pipette, 90 µL of YLG or dYLG solution was added to triplicate wells, and the plate immediately loaded into a BioTek Synergy HTX plate reader. The program parameters included incubation at 25 °C, an initial shaking step at 425 rpm for 20 seconds, followed by fluorescence detection with excitation/emission of 485/520 nm (top optics, gain 35, read height of 4.5 mm). Readings were taken every 2 minutes over a span of 30 minutes. The hydrolysis rate was determined by assessing the change in fluorescence over the initial 10 minutes, post-subtraction of background fluorescence observed in no-protein control samples. K_d_ values were derived in GraphPad Prism, applying the in-built ‘Michaelis-Menten’ model. Results were displayed with 95% confidence interval values.

### Intrinsic tryptophan fluorescence (ITF)

ITF assays were performed in 384-well black microplates (Greiner 781076) with four technical replicates of 20 µL reactions comprising 10 µM protein, 20 mM HEPES pH 7.5, 150 mM NaCl, 5% [v/v] DMSO, and 0–400 µM ligand. Ligands were diluted 10-fold into ITF buffer (20 mM HEPES, 150 mM NaCl, pH 7.5) from 20× stocks in DMSO, and dispensed at 10 µL per well with a multichannel pipette. Protein was diluted to 2× concentration in ITF buffer and dispensed at 10 µL per well with a repeat pipettor. The plate was mixed at 120 rpm for 2 min, centrifuged at 500×*g* for 2 min and incubated in the dark for 30 minutes at ambient temperature. Protein fluorescence was measured at 25 °C on a CLARIOstar multimode plate reader (BMG Labtech), with excitation at 295 ± 10 nm, emission at 360 ± 10 nm, and a 325 nm longpass dichroic filter. Each measurement consisted of 20 flashes per sample. Ligand-only samples were used as blank. Data was plotted in GraphPad Prism as non-linear regression and the in-built ‘[Inhibitor] vs response (three parameters)’ model with IC_50_ > 0.

### β-galactosidase reporter assays

Cell pellets were resuspended in 500 µL Z-buffer (60 mM Na_2_HPO_4_, 40 mM NaH_2_PO_4_, 10 mM KCl, 1 mM MgSO_4_, 50 mM β-mercaptoethanol, pH 7.0), 50 µL chloroform, 25 µL 0.1% [w/v] SDS. The samples were lysed by vortex-mixing for 10 seconds followed by incubation at 28 °C for five minutes. The reaction was initiated by adding 100 µL o-nitrophenyl-β-D-galactopyranoside (4 mg/mL), and allowed to run for 1 h at 28 °C. The reaction was terminated by adding 100 µL of 2.5 M Na_2_CO_3_, and samples were centrifuged at 12,000×*g* for five minutes to remove cell debris. The hydrolysis of o-nitrophenyl-β-D-galactopyranoside was measured by loading 150 µL of clarified supernatant into a 96-well, clear microtitre plate (Greiner 655095), and reading the absorbance at 420 nm using a Synergy HTX plate reader (BioTek). Enzyme activity (expressed as Miller units, average of three replicates) was calculated with the formula: (OD_420_ × 1000) / (time of reaction [min] × volume on plate [mL] × OD_600_).

### Gene expression analysis by RT–qPCR

RNA extraction, DNase treatment, and cDNA synthesis were performed following the method described previously^6^ with slight modification of using small glass beads 212-300 μm (50-70 U.S. sieve, Sigma) for grinding. Quantitative RT-PCR was conducted on a Roche Lightcycler 480 instrument using Luna Universal qPCR master mix (New England Biolabs in Ipswich, MA, USA). The geometric mean of *recA* and *gyrB* transcripts was used to normalise data. Target genes were amplified using the primers listed in Table S6. All RT–qPCR analyses were performed in technical duplicates using three independently isolated RNA samples (biological triplicates).

### Total transcriptome analysis

RNA was extracted followed by DNase I treatment. Only samples with RNA integrity scores ≥ 7, as determined by an Agilent 2200 Tape Station, were used for library preparation. The libraries were generated using Illumina Stranded Ribo-Zero and a ribodepletion whole-transcriptome workflow as per the manufacturer instruction. Libraries were sequenced on our NovaSeq 6000 instrument and across one lane of an SP-300 cycle flow cell (150PE) with v1.5 chemistry. Adapter trimming, read mapping onto the *Bacillus subtilis* subsp. *subtilis* str. 168 (GCA_000009045) reference genome and read counts were carried out with kallisto^54^. Differentially expressed gene (DEG) analysis was done by DEseq2 ^55^ using R studio. For each transcript, a false-discovery rate-adjusted P value1<10.05 and log_2_ fold change (FC) >1 or <−1 were set as significance thresholds. Venn diagrams were generated using a web-based tool (https://bioinformatics.psb.ugent.be/webtools/Venn/; Bioinformatics & Evolutionary Genomics Technologiepark, Gent, Belgium). Volcano plots were generated with R package enhancedvolcano; heatmap and DEG clustering were generated using R package pheatmap in R Studio.

### Biofilm quantification with crystal violet

Biofilm quantification used a modified version of a procedure described previously^56^. After adequate biofilm growth, the liquid medium was gently removed from each well, and the well rinsed once with 2 mL of water. The biofilm was then fixed by incubating at 60 °C for one hour. Each well was stained with 3 mL 0.1% [w/v] crystal violet solution (dissolved in 75% methanol) for 30 minutes. After staining, wells were washed three times with water. The crystal violet stain bound to the biofilm was eluted with 3 mL of 30% [v/v] acetic acid. The intensity of the stain, which correlates with biofilm biomass, was quantified by measuring the absorbance at 575 nm using a BioTek Synergy HTX plate reader and an appropriate dilution factor to ensure signal linearity.

## SUPPLEMENTARY INFORMATION

### Supplementary Figures

**Figure S1.**
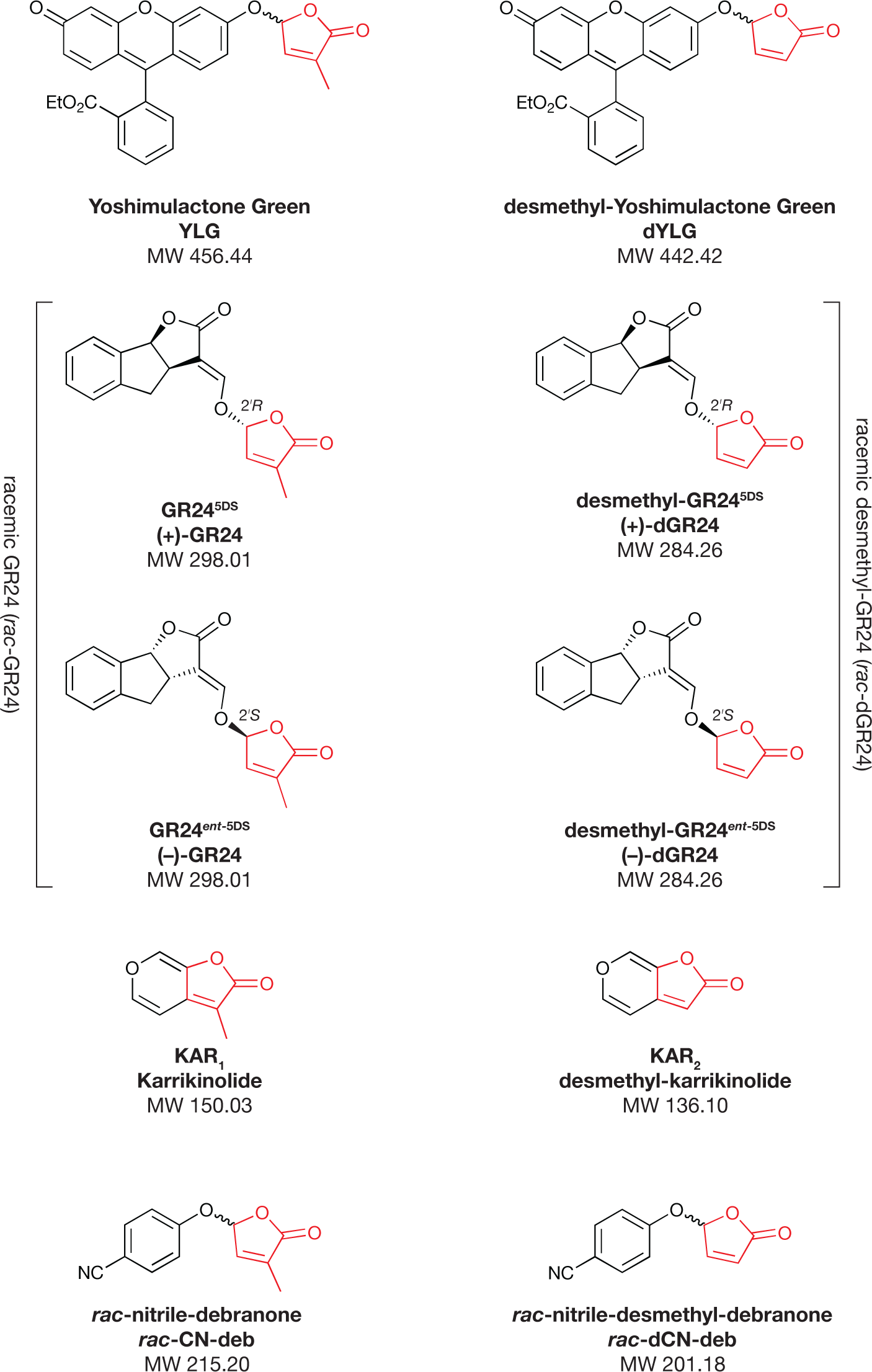
Chemical structures, abbreviations and synonyms of compounds used in this study. Compounds with a methyl-substituted butenolide moiety (red) are aligned on the left; desmethyl butenolide compounds on the right. Where appropriate, the stereochemistry at the 2ʹ carbon of the butenolide ring is indicated. Note that YLG and dYLG are used as racemates.

**Figure S2.**
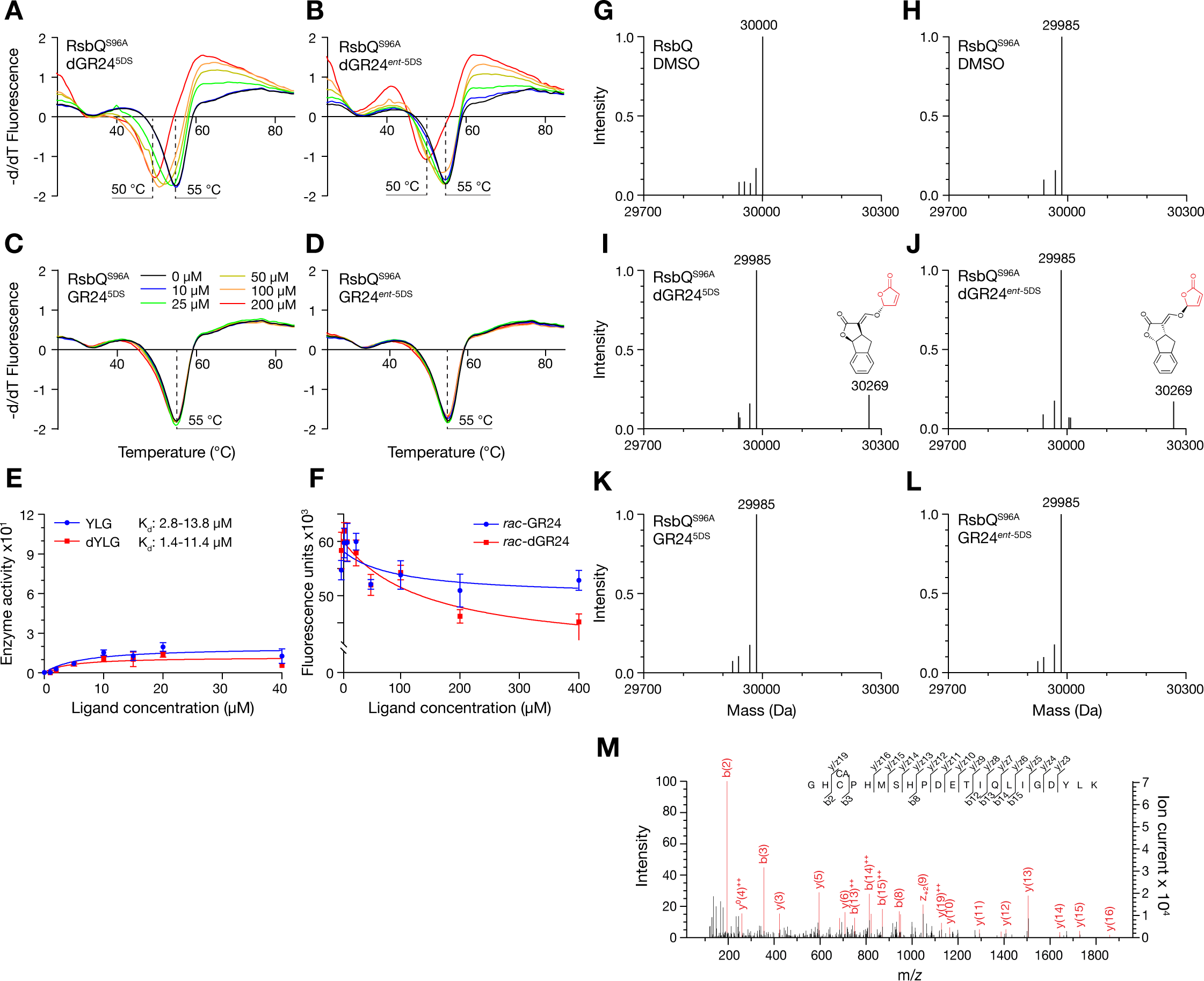
Serine 96 is necessary for functional interaction of RsbQ with butenolides. **A–D**. Thermal shift assays of purified SUMO-RsbQ^S96A^ mutant fusion protein challenged with increasing concentrations of the two constituent enantiomers of desmethyl-GR24 (dGR24^5DS^ and dGR24*^ent^*^-5DS^) or the two corresponding enantiomers of GR24 (C, D). Inferred protein melting temperatures are indicated with dashed lines. **E**. Hydrolysis activity of SUMO-RsbQ^S96A^ towards the profluorescent butenolides Yoshimulactone Green (YLG) and desmethyl-YLG (dYLG). Data are means ± SD of n=3 technical replicates. K_d_ values were estimated by non-linear regression and are given as 95% confidence intervals. **F**. Intrinsic tryptophan fluorescence measurements of SUMO-RsbQ^S96A^ in response to varying concentrations of racemic GR24 or dGR24. Data are means ± SD of n=4 technical replicates. **G,H**. Deconvoluted intact mass spectra of purified wild type RsbQ (calculated MW=30,000 Da) and RsbQ^S96A^ mutant (calculated MW=29,984 Da) treated with DMSO alone as negative controls. **I–L**. Deconvoluted intact mass spectra of purified wild type RsbQ^S96A^ mutant protein treated with individual enantiomers of dGR24 and GR24. Shifted peaks are labelled with observed MW and the chemical structure of putative adducts consistent with the treatment and increase in mass. **M**. Tandem mass spectrum of a fragmented peptide of RsbQ (Gly246 to Lys266; M_mi_ = 2390.1 Da) following incubation with DMSO and trypsin digestion. The heavier peptide (M_mi_ = 2472.1 Da) that was modified with a putative desmethyl butenolide and isolated from wild type RsbQ was not detected in any experiment with DMSO. The peptide was identified using MASCOT (version 2.5.1, Matrix Sciences) with ions (red peaks) supporting identification of the peptide sequence shown. Identified peaks representing c/b and y/z ions are annotated with ^++^, indicating a doubly-charged ion; ^0^ representing an ion − H_2_O and * an ion − NH_3_; CA, carbamidomethyl, a cysteine modification caused by iodoacetamide.

**Figure S3.**
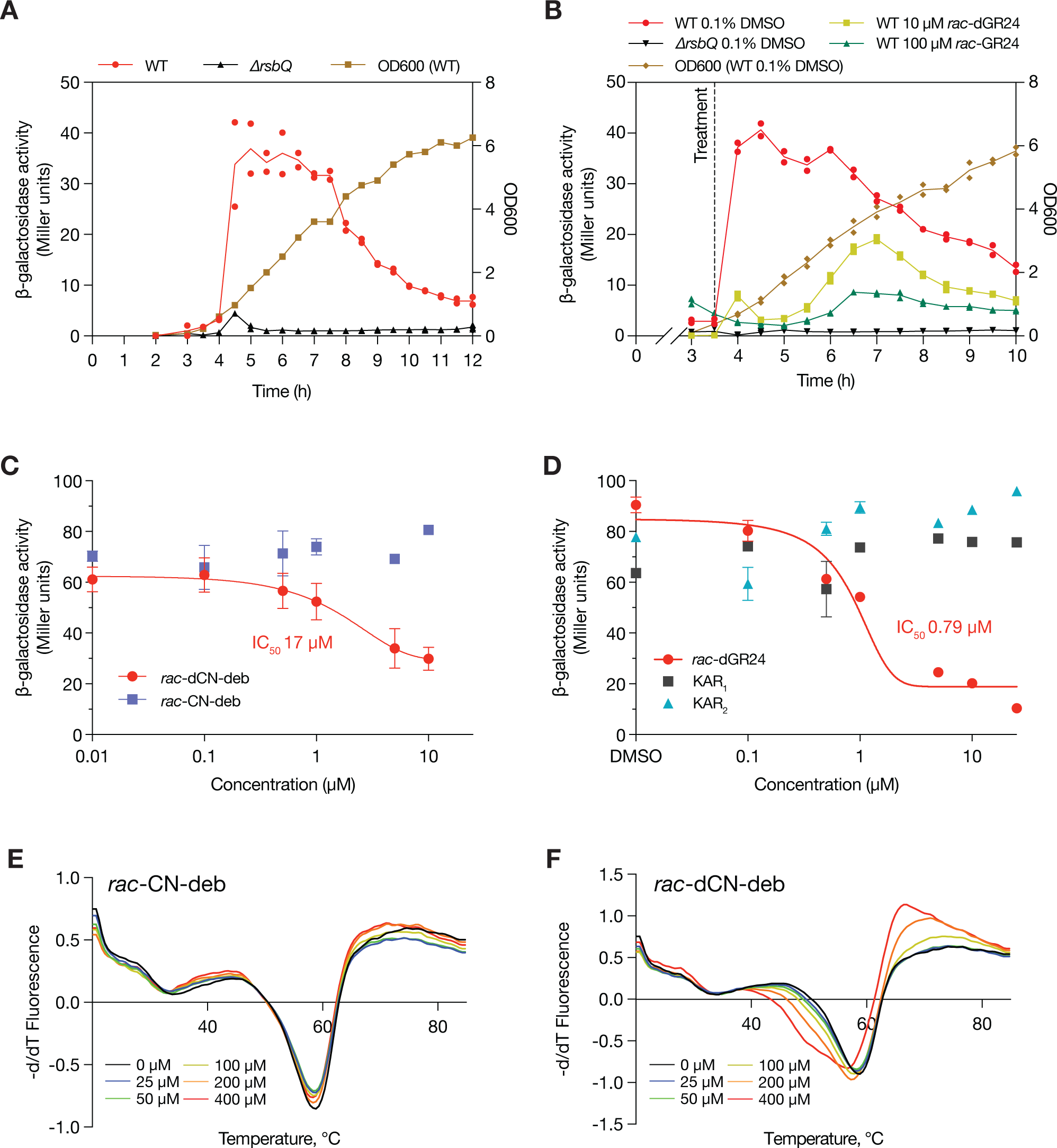
Characterisation of RsbQ-mediated responses to karrikins and debranones. **A.** Natural induction of β-galactosidase activity from the *amyE::ctc-LacZ* reporter in WT *B. subtilis* 168 (strain PB198) and an isogenic *ΔrsbQ* null mutant (strain PB605) as the culture transitions from exponential into stationary phase. Cultures (50 mL) were inoculated at time zero with a 1:1000 dilution of an overnight starter culture, grown at 37 °C with shaking at 200 RPM. Data show individual values from two experimental replicates. **B.** Effects of 100 µM *rac*-GR24 and 10 µM *rac*-dGR24 on induction of β-galactosidase activity. Note the different concentrations. Vertical dashed line indicates addition of indicated chemical treatment. Cultures were grown as per **A**. Data show individual values from two experimental replicates. **C, D**. Response of WT *B. subtilis* 168 strain PB198 to increasing concentrations of (**C**) *rac*-nitrile-debranone (CN-deb) or *rac-*desmethyl-nitrile-debranone (dCN-deb), and (**D**) karrikins KAR_1_ and KAR_2_. *rac*-dGR24 is included as a positive control. Data are means ± SE of n=3 replicate samples from a single treated culture for each concentration/compound. Cultures (50 mL) were grown at 30 °C and shaken at 200 rpm; treatment commenced at OD_600_ of 0.5, and samples were harvested 2 hours later. **E, F**. Thermal shift assays of purified SUMO-RsbQ fusion protein challenged with 0–400 µM *rac*-CN-deb (**E**) or *rac*-dCN-deb (**F**). Only a weak shift in protein melt profile is observed with >200 µM *rac*-dCN-deb.

**Figure S4.**
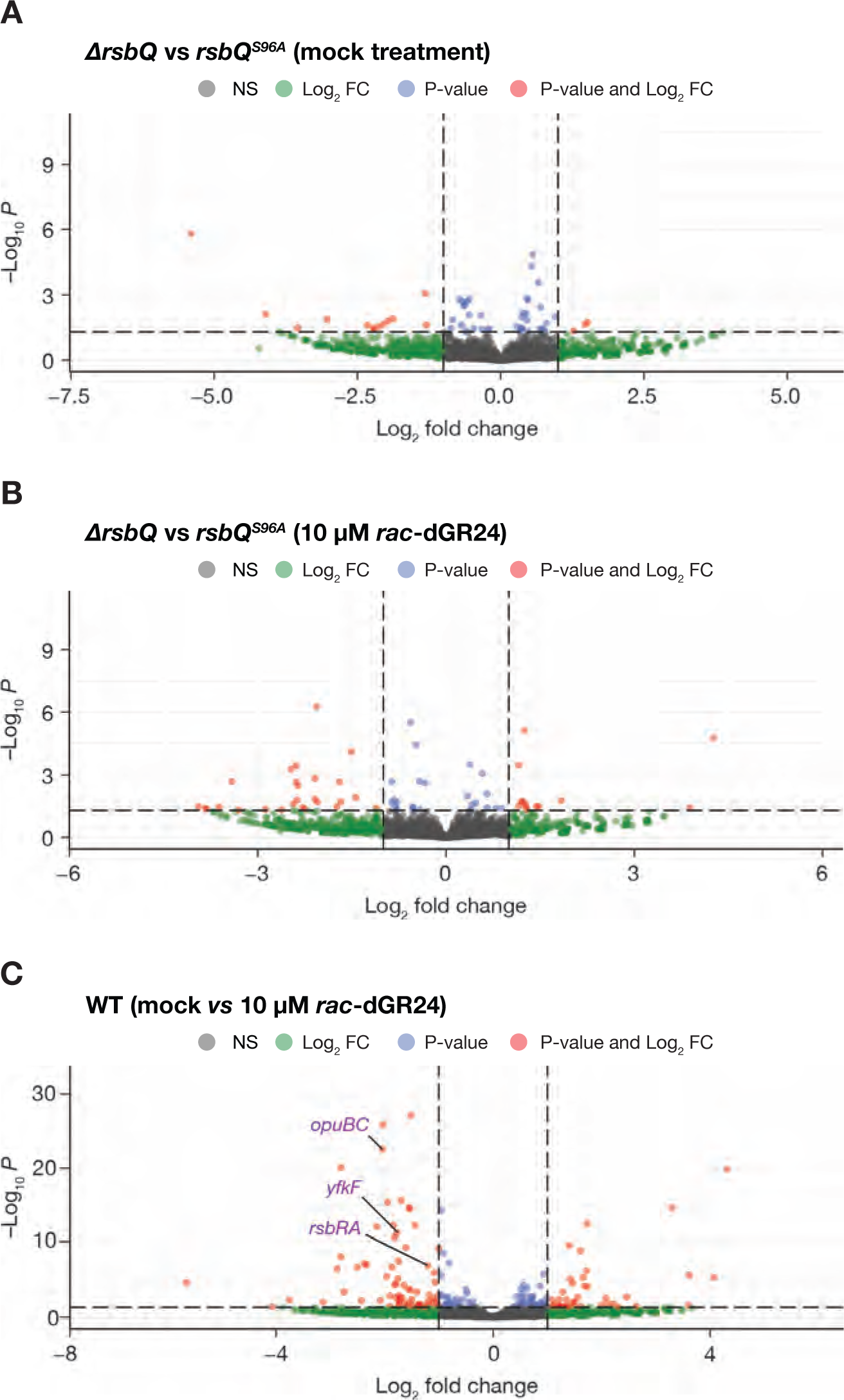
The *ΔrsbQ* and *rsbQ^S96A^* mutants respond differently to *rac*-dGR24. Volcano plots highlighting differentially expressed genes (DEGs) following RNA-seq. Those that passed the criteria of log_2_fold change >1 or <−1, and false discovery rate-adjusted P-value < 0.05 (thresholds indicated by dashed lines), are plotted as red circles. A. Comparison of *ΔrsbQ* and *rsbQ^S96A^* mutants mock-treated with 0.1% DMSO. A total of 16 genes were defined as differentially expressed under these conditions. B. Comparison of *ΔrsbQ* and *rsbQ^S96A^* mutants treated with 10 µM *rac*-dGR24. A total of 39 genes were defined as differentially expressed under these conditions. C. Comparison of mock-treated and *rac*-dGR24-treated WT transcriptomes, identifying a total of 109 DEGs. As an example, three DEGs from Figure 3F that are σ^B^-dependent and repressed by *rac*-dGR24 are labelled.

**Figure S5.**
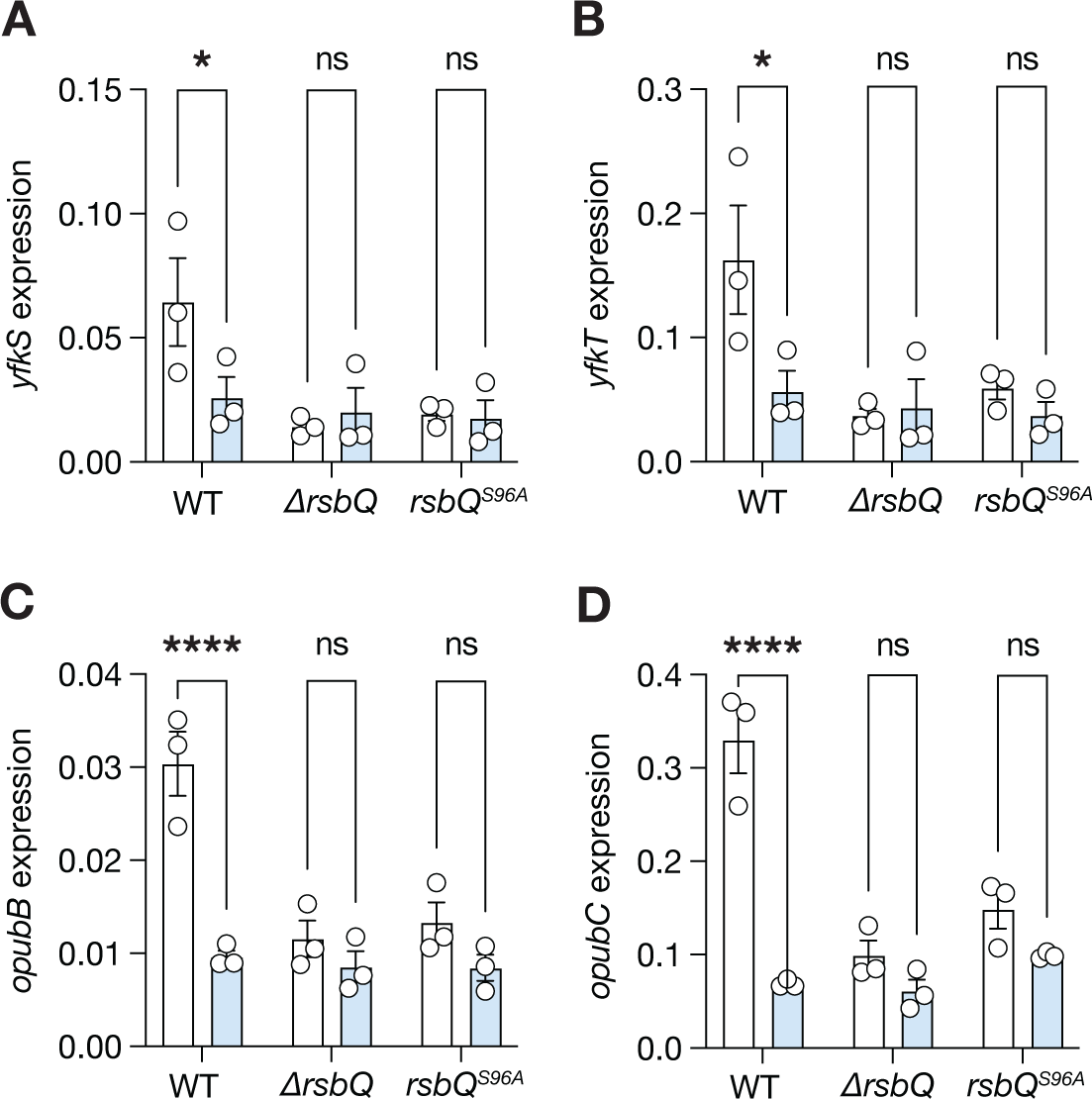
Responses of RsbQ-dependent transcripts to *rac*-dGR24. Levels of selected transcripts following treatment with racemic dGR24, determined by qRT-PCR. Cultures were treated and harvested as described in Figure 3. All transcripts were normalised to *gyrB* and *recA* reference transcripts. Data are means ± SE of n=3 replicate cultures. Significant differences for pairwise comparisons within a genotype are indicated: * P<0.05; **** P<0.0001; ns, non-significant (two-way ANOVA).

**Figure S6.**
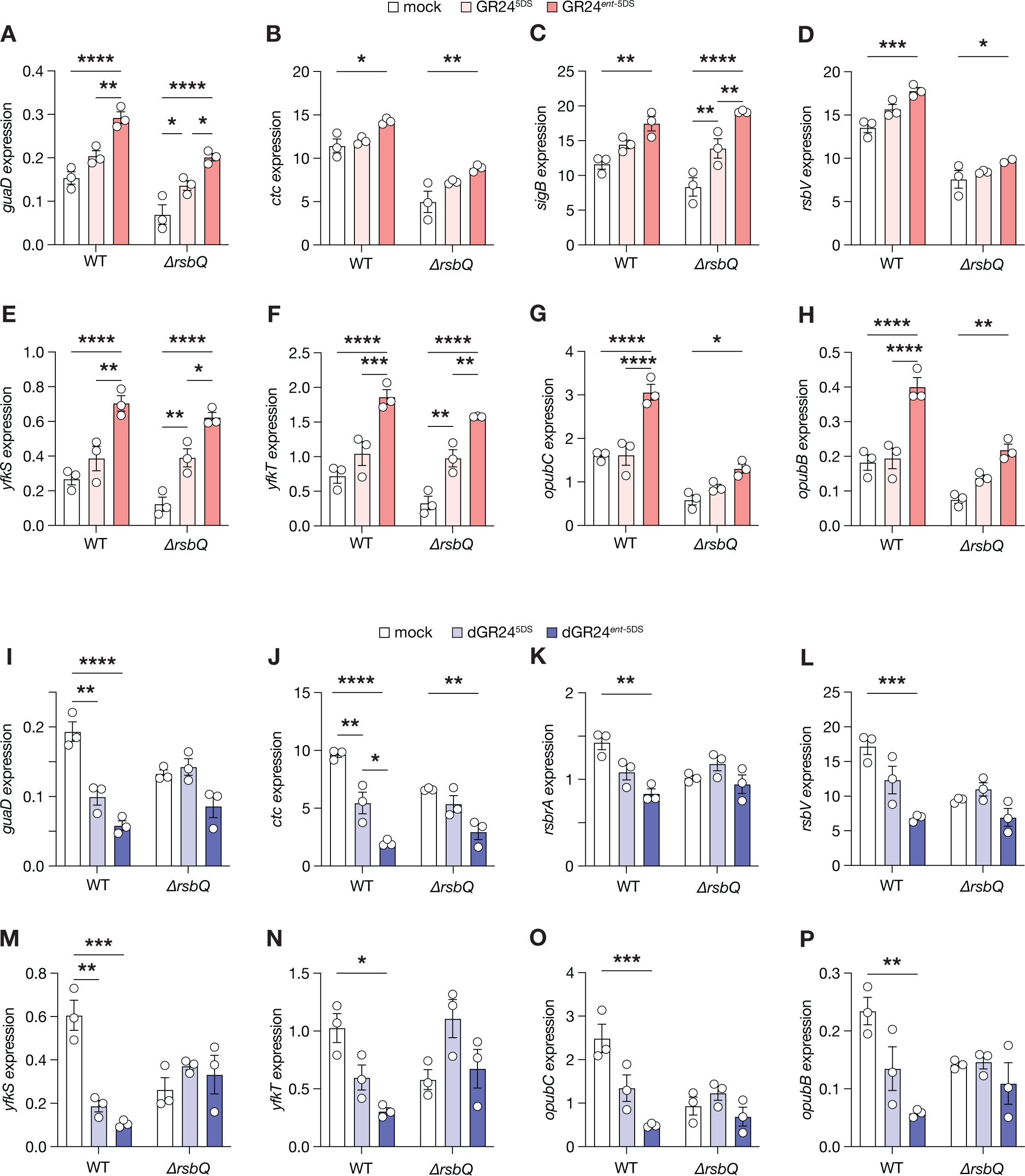
Responses of σ^B^-dependent transcripts to individual enantiomers of GR24 and dGR24. Levels of selected transcripts following treatment with individual enantiomers of GR24 (A–H) or dGR24 (I–P), determined by qRT-PCR. Cultures were treated and harvested as described in Figure 3. All transcripts were normalised to *gyrB* and *recA* reference transcripts. Data are means ± SE of n=3 replicate cultures. Significant differences for pairwise comparisons within a genotype are indicated: * P<0.05; ** P<0.01; *** P<0.001; **** P<0.0001; for clarity, non-significant comparisons are omitted (two-way ANOVA).

### Supplementary Datasets

**Data S1** RsbQ, KAI2 and D14 homologues used for conservation analysis of pocket residues.

**Data S2** Alignment of RsbQ sequences used in Figure 1.

**Data S3** Alignment of KAI2 sequences used in Figure 1.

**Data S4** Alignment of D14 sequences used in Figure 1.

## KEY RESOURCES TABLE

The table highlights the reagents, genetically modified organisms and strains, cell lines, software, instrumentation, and source data **essential** to reproduce results presented in the manuscript. Depending on the nature of the study, this may include standard laboratory materials (i.e., food chow for metabolism studies, support material for catalysis studies), but the table is **not** meant to be a comprehensive list of all materials and resources used (e.g., essential chemicals such as standard solvents, SDS, sucrose, or standard culture media do not need to be listed in the table). **Items in the table must also be reported in the method details section within the context of their use.** To maximize readability, the number of **oligonucleotides and RNA sequences** that may be listed in the table is restricted to no more than 10 each. If there are more than 10 oligonucleotides or RNA sequences to report, please provide this information as a supplementary document and reference the file (e.g., See Table S1 for XX) in the key resources table.

***Please note that ALL references cited in the key resources table must be included in the main references list.*** Please report the information as follows:

- **REAGENT or RESOURCE:** Provide the full descriptive name of the item so that it can be identified and linked with its description in the manuscript (e.g., provide version number for software, host source for antibody, strain name). In the experimental models section (applicable only to experimental life science studies), please include all models used in the paper and describe each line/strain as: model organism: name used for strain/line in paper: genotype. (i.e., Mouse: OXTR^fl/fl^: B6.129(SJL)-Oxtr^tm1.1Wsy/J^). In the biological samples section (applicable only to experimental life science studies), please list all samples obtained from commercial sources or biological repositories. Please note that software mentioned in the methods details or data and code availability section needs to also be included in the table. See the sample tables at the end of this document for examples of how to report reagents.
- **SOURCE:** Report the company, manufacturer, or individual that provided the item or where the item can be obtained (e.g., stock center or repository). For materials distributed by Addgene, please cite the article describing the plasmid and include “Addgene” as part of the identifier. If an item is from another lab, please include the name of the principal investigator and a citation if it has been previously published. If the material is being reported for the first time in the current paper, please indicate as “this paper.” For software, please provide the company name if it is commercially available or cite the paper in which it has been initially described.
- **IDENTIFIER:** Include catalog numbers (entered in the column as “Cat#” followed by the number, e.g., Cat#3879S). Where available, please include unique entities such as <u>RRIDs</u>, Model Organism Database numbers, accession numbers, and PDB, CAS, or CCDC IDs. For antibodies, if applicable and available, please also include the lot number or clone identity. For software or data resources, please include the URL where the resource can be downloaded. Please ensure accuracy of the identifiers, as they are essential for generation of hyperlinks to external sources when available. Please see the Elsevier <u>list of data repositories</u> with automated bidirectional linking for details. When listing more than one identifier for the same item, use semicolons to separate them (e.g., Cat#3879S; RRID: AB_2255011). If an identifier is not available, please enter “N/A” in the column.

`o` ***A NOTE ABOUT RRIDs:*** We highly recommend using RRIDs as the identifier (in particular for antibodies and organisms but also for software tools and databases). For more details on how to obtain or generate an RRID for existing or newly generated resources, please <u>visit the RII</u> or <u>search for RRIDs</u>.

Please use the empty table that follows to organize the information in the sections defined by the subheading, skipping sections not relevant to your study. Please do not add subheadings. To add a row, place the cursor at the end of the row above where you would like to add the row, just outside the right border of the table. Then press the ENTER key to add the row. Please delete empty rows. Each entry must be on a separate row; do not list multiple items in a single table cell. Please see the sample tables at the end of this document for relevant examples in the life and physical sciences of how reagents and instrumentation should be cited.

### TABLE FOR AUTHOR TO COMPLETE

Please upload the completed table as a separate document. **<u>Please do not add subheadings to the key resources</u> <u>table.</u>** If you wish to make an entry that does not fall into one of the subheadings below, please contact your handling editor. **<u>Any subheadings not relevant to your study can be skipped.</u>** (**NOTE:** References within the KRT should be in numbered style rather than Harvard.)

#### Key resources table

| REAGENT or RESOURCE | SOURCE | IDENTIFIER |
| --- | --- | --- |
| Bacterial and virus strains |  |  |
| *amyE::ctc-lacZ, trpC2* | Brody 2001 | PB198 |
| Δ*rsbQ, amyE::ctc-lacZ, trpC2* | Brody 2001 | PB605 |
| *rsbQ*<sup>S96A</sup>, *amyE::ctc-lacZ, trpC2* | Brody 2001 | PB634 |
| NCIB 3610 *ComI*(Q12L) | Konkol 2013 | DK1042 |
| NCIB 3610 *ComI*(Q12L), Δ*rsbQP::cm* | This study | WLB042 |
| Biological samples |  |  |
| Chemicals, peptides, and recombinant proteins |  |  |
| Deposited data |  |  |
| Raw and analysed data | This paper | GSE236759 |
| Experimental models: Organisms/strains |  |  |
| Oligonucleotides |  |  |
| Recombinant DNA |  |  |
| Software and algorithms |  |  |
| Kallisto | Bray et al.<sup>1</sup> | https://github.com/pachterlab/kallisto |
| R Studio 4.2.3 | RStudio | https://www.rstudio.com/ |
| DEseq2 version 4.3 | Love et al.<sup>2</sup> | https://bioconductor.org/packages/release/bioc/html/DESeq2.html |
| Venn diagrams | Bioinformatics & Evolutionary Genomics<br>Technologiepark 927<br>B-9052 Gent<br>BELGIUM | https://bioinformatics.psb.ugent.be/webtools/Venn/ |
| R package pheatmap version 1.0.12 | Raivo Kolde | https://cran.r-project.org/web/packages/pheatmap/index.html |
| R package EnhancedVolcano version 4.3 | Blighe et al.<sup>3</sup> | https://github.com/kevinblighe/EnhancedVolcano |
| Other |  |  |

### LIFE SCIENCE TABLE WITH EXAMPLES FOR AUTHOR REFERENCE

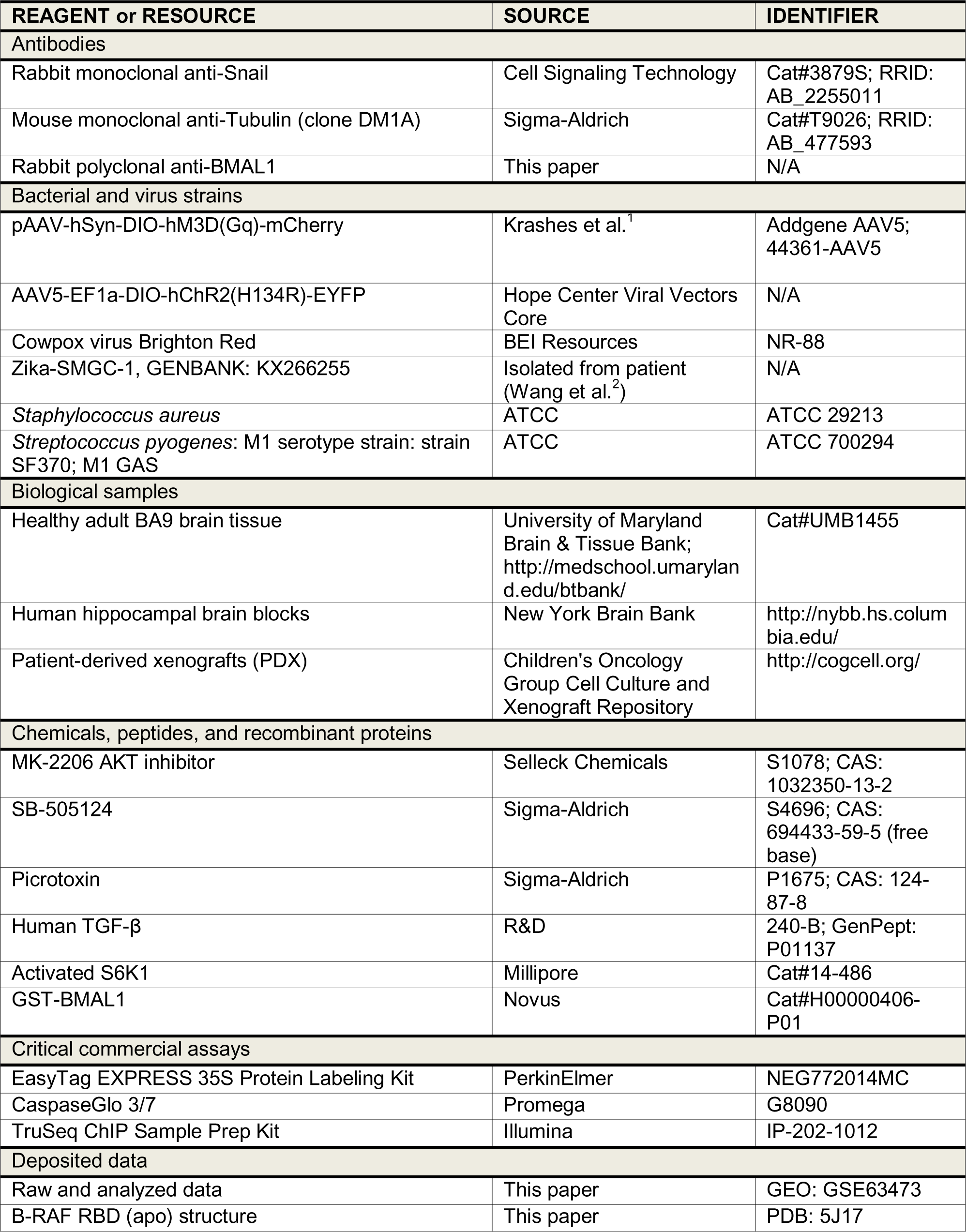
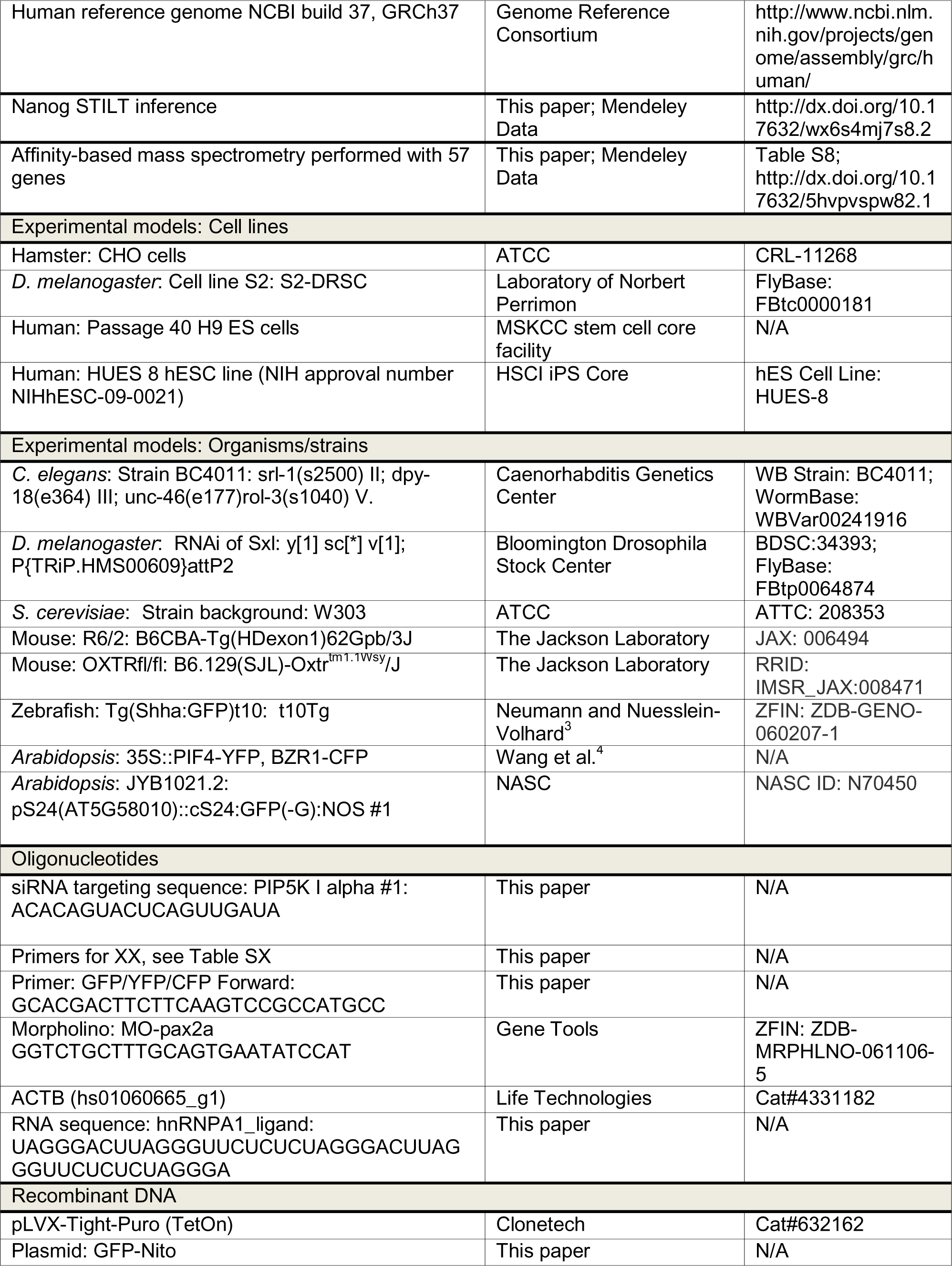
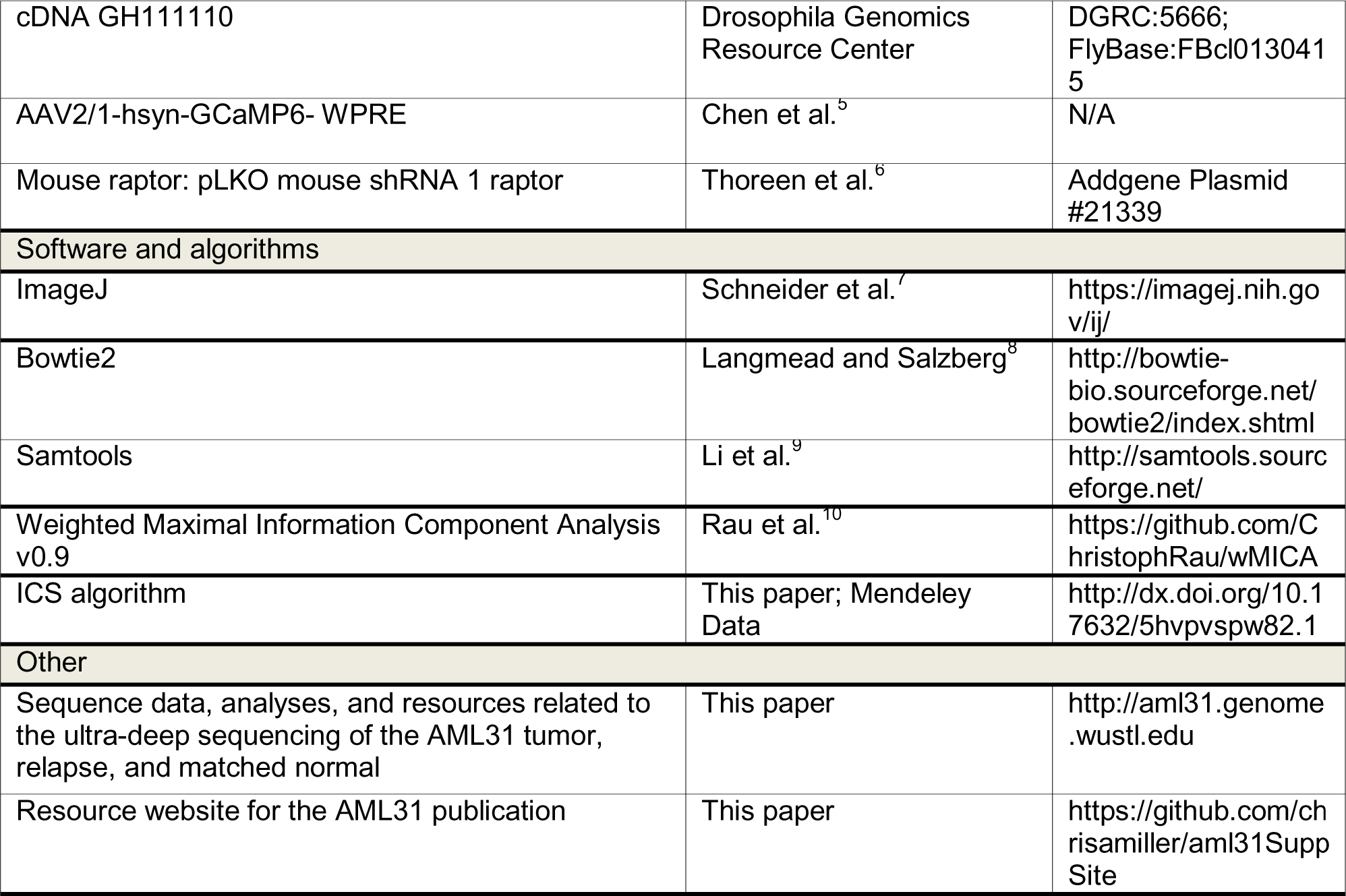

### PHYSICAL SCIENCE TABLE WITH EXAMPLES FOR AUTHOR REFERENCE

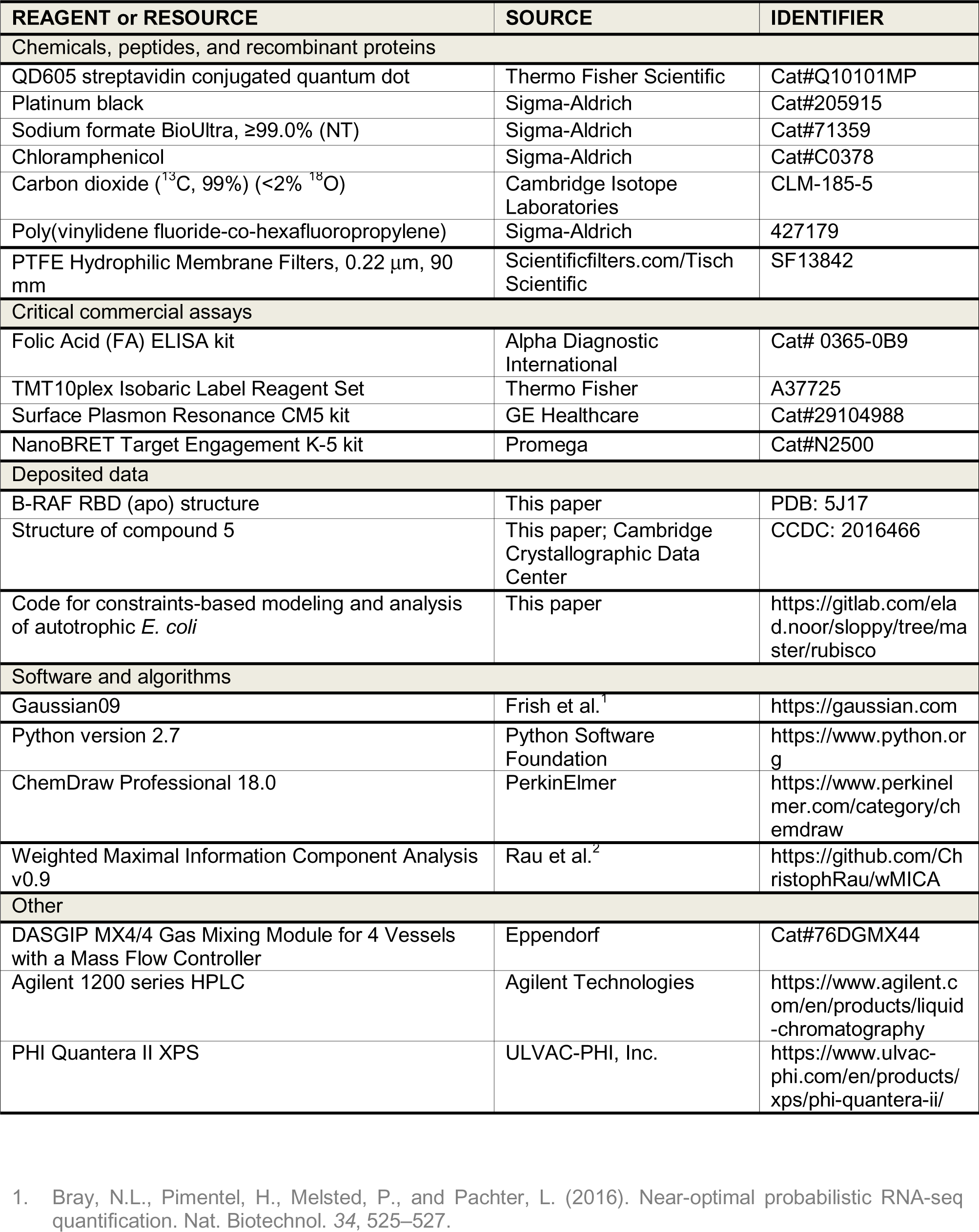
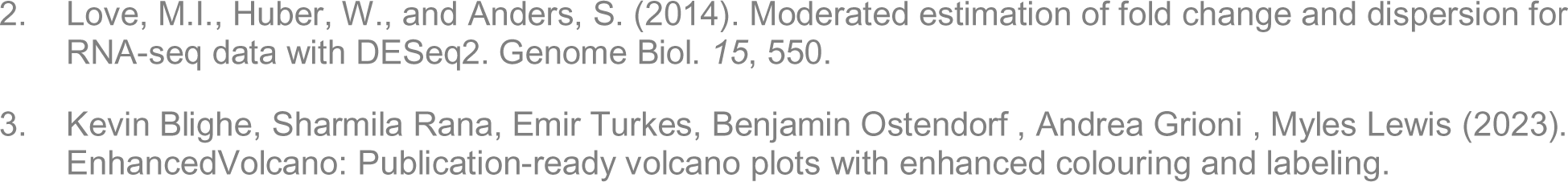

| REAGENT or RESOURCE | SOURCE | IDENTIFIER |
| --- | --- | --- |
| Chemicals, peptides, and recombinant proteins |  |  |
| QD605 streptavidin conjugated quantum dot | Thermo Fisher Scientific | Cat#Q10101MP |
| Platinum black | Sigma-Aldrich | Cat#205915 |
| Sodium formate BioUltra, ≥99.0% (NT) | Sigma-Aldrich | Cat#71359 |
| Chloramphenicol | Sigma-Aldrich | Cat#C0378 |
| Carbon dioxide (<sup>13</sup>C, 99%) (<2% <sup>18</sup>O) | Cambridge Isotope Laboratories | CLM-185-5 |
| Poly(vinylidene fluoride-co-hexafluoropropylene) | Sigma-Aldrich | 427179 |
| PTFE Hydrophilic Membrane Filters, 0.22 μm, 90 mm | Scientificfilters.com/Tisch Scientific | SF13842 |
| Critical commercial assays |  |  |
| Folic Acid (FA) ELISA kit | Alpha Diagnostic International | Cat# 0365-0B9 |
| TMT10plex Isobaric Label Reagent Set | Thermo Fisher | A37725 |
| Surface Plasmon Resonance CM5 kit | GE Healthcare | Cat#29104988 |
| NanoBRET Target Engagement K-5 kit | Promega | Cat#N2500 |
| Deposited data |  |  |
| B-RAF RBD (apo) structure | This paper | PDB: 5J17 |
| Structure of compound 5 | This paper; Cambridge Crystallographic Data Center | CCDC: 2016466 |
| Code for constraints-based modeling and analysis of autotrophic *E. coli* | This paper | https://gitlab.com/elad.noor/sloppy/tree/master/rubisco |
| Software and algorithms |  |  |
| Gaussian09 | Frish et al.<sup>1</sup> | https://gaussian.com |
| Python version 2.7 | Python Software Foundation | https://www.python.org |
| ChemDraw Professional 18.0 | PerkinElmer | https://www.perkinelmer.com/category/chemdraw |
| Weighted Maximal Information Component Analysis v0.9 | Rau et al.<sup>2</sup> | https://github.com/ChristophRau/wMICA |
| Other |  |  |
| DASGIP MX4/4 Gas Mixing Module for 4 Vessels with a Mass Flow Controller | Eppendorf | Cat#76DGMX44 |
| Agilent 1200 series HPLC | Agilent Technologies | https://www.agilent.com/en/products/liquid-chromatography |
| PHI Quantera II XPS | ULVAC-PHI, Inc. | https://www.ulvac-phi.com/en/products/xps/phi-quantera-ii/ |

1. Bray, N.L., Pimentel, H., Melsted, P., and Pachter, L. (2016). Near-optimal probabilistic RNA-seq quantification. *Nat. Biotechnol.* 34, 525–527.
2. Love, M.I., Huber, W., and Anders, S. (2014). Moderated estimation of fold change and dispersion for RNA-seq data with DESeq2. *Genome Biol.* 15, 550. 3. Kevin Blighe, Sharmila Rana, Emir Turkes, Benjamin Ostendorf , Andrea Grioni , Myles Lewis (2023). EnhancedVolcano: Publication-ready volcano plots with enhanced colouring and labeling.

